# Conserved structures of neural activity in sensorimotor cortex of freely moving rats allow cross-subject decoding

**DOI:** 10.1101/2021.03.04.433869

**Authors:** Svenja Melbaum, Eleonora Russo, David Eriksson, Artur Schneider, Daniel Durstewitz, Thomas Brox, Ilka Diester

## Abstract

Our knowledge about neuronal activity in the sensorimotor cortex relies primarily on stereotyped movements that are strictly controlled in experimental settings. It remains unclear how results can be carried over to less constrained behavior like that of freely moving subjects. Toward this goal, we developed a self-paced behavioral paradigm that encouraged rats to engage in different movement types. We employed bilateral electrophysiological recordings across the entire sensorimotor cortex and simultaneous paw tracking. These techniques revealed behavioral coupling of neurons with lateralization and an anterior–posterior gradient from the premotor to the primary sensory cortex. The structure of population activity patterns was conserved across animals despite the severe under-sampling of the total number of neurons and variations in electrode positions across individuals. We demonstrated cross-subject and cross-session generalization in a decoding task through alignments of low-dimensional neural manifolds, providing evidence of a conserved neuronal code

**One-sentence summary:** Similarities in neural population structures across the sensorimotor cortex enable generalization across animals in the decoding of unconstrained behavior.

**Graphical abstract:** Conserved structures of neural activity in freely moving rats allow for cross-subject decoding.
(a) We conducted electrophysiological recordings across the bilateral sensorimotor cortex of six freely moving rats. Neural activities were projected into a low-dimensional space with LEMs (*22*). (b) In a decoding task, points in the aligned low-dimensional neural state space were used as input for a classifier that predicted behavioral labels. Importantly, training and testing data originated from different rats. (c) Our procedure led to successful cross-subject generalization for sessions with sufficient numbers of recorded units. The rat and brain drawings are adapted from `scalablebrainatlas.incf.org` and SciDraw.

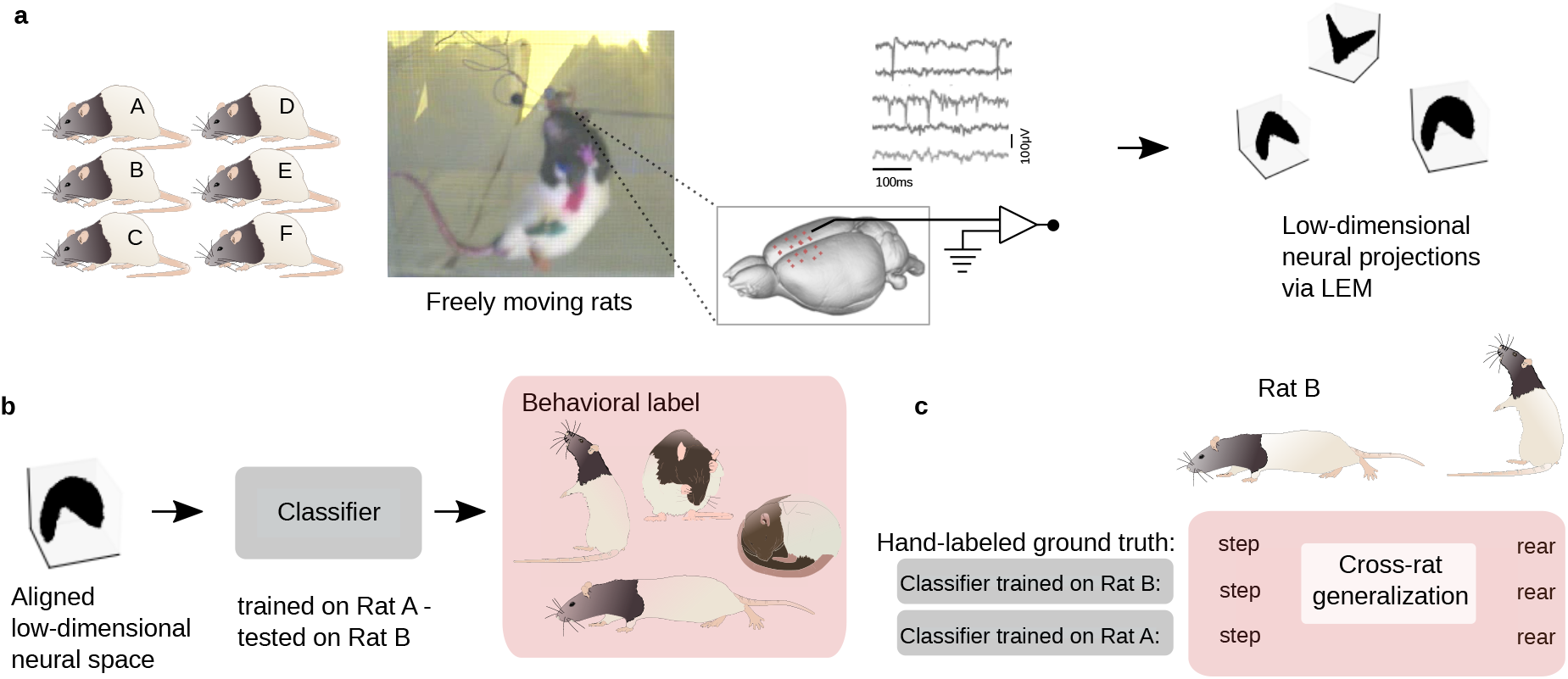

## Introduction

Humans and animals are capable of generating a vast array of behaviors. This feature is dependent on the brain’s ability to generate a wide repertoire of neural activity patterns, which may rely on subsets of general motifs (*1*). Experimental, computational, and theoretical work has identified the rich underlying structures within neural populations regarding movement control, decision-making, and memory tasks (*2*). Similarities in population structures across different modalities such as speech and arm movements (*3*), as well as the relevance of population-level phenomena to learning (*4*), hint at the existence of general principles that could be shared across subjects. For simple, constrained behavior such as running on a linear track, population structures in some brain regions such as the hippocampus seem to be conserved, even across subjects (*5*). Similarities in neural population structures have not yet been shown for freely roaming animals and various naturally occurring behaviors. Whether population structures are sufficiently conserved across subjects to allow for the cross-subject decoding of behavioral categories remains an open question in systems neuroscience. This question has great implications for neuro-prosthetic approaches, among other research topics. Such conservation of neural structures would allow for a shorter adaptation or fine-tuning phase of Brain-Machine-Interface (BCI) systems from one subject to another as opposed to training the system from scratch.

We addressed this question with non-linear mapping applied to electrophysiological recordings across the entire bilateral sensorimotor cortex of the rat. The neural trajectories of dynamical systems have been suggested as a method to understand neural activity (*6, 7, 8, 9, 10, 11, 12, 13, 14, 15, 16, 4, 17, 18, 19, 20, 21*). Therefore, we built on Laplacian Eigenmaps (LEMs) (*22, 5*), which map high-dimensional data via the data’s affinity to a low-dimensional manifold. When affinities are defined according to neuronal population activity, they can be used as tools to visualize structures and relationships among population activities at different time points of a recording session in a low-dimensional space. This can potentially reveal conserved structures across sessions and animals (*5*).

To investigate the degree to which low-dimensional structures are conserved, it is necessary to involve several types of behavior. In principle, it is possible to train animals in different tasks, but this has several limitations: (1) training animals is time-consuming, especially if multiple behaviors are involved; (2) the trained behavior often results in stereotyped movements and (due to the plasticity of the mammalian brain) corresponding changes in neuronal representations; and (3) frequent transitions between behaviors are not feasible. Furthermore, spontaneous movements influence neuronal activity, even in well-controlled tasks (*23*). Therefore, we refrained from controlling the behavior from the start, instead allowing the rats to roam freely in a Plexiglas box. Consequently, the animals showed their full array of natural behavior, such as rearing, grooming, turning, stepping, drinking, and resting, in an unbiased manner.

To verify this approach, we first compared neuronal activity with previously reported results from more constrained behaviors by focusing on step- and swing-like paw movements. This study confirmed that the quality of information conveyed by our recorded data was comparable to that found in conventionally controlled settings. In addition, we reported a strong anterior– posterior gradient in the lateralization of forelimb representations from the premotor cortex to the primary sensory cortex. This gradient emphasizes the strong involvement of more posterior regions in the encoding of step-like behavior.

After this validation, we focused on analyzing the population code for more complex behaviors. We conducted a normal within-session decoding experiment to show that the neuronal code contains enough information about the behavior class. Across sessions, the signals of individual neurons were not comparable since neurons typically cannot be traced over multiple days. Across subjects, even the electrode positions varied. However, we found evidence that the signal from the *population* of neurons shared a common structure across sessions and even across subjects. In particular, decoding behavioral categories from the neuronal population activity was possible across different subjects.

## Results

Rats moved unconstrained in a rectangular arena and conducted movements in different behavioral categories (i.e., stepping, turning, drinking, grooming, and rearing) while searching for water drops, which a robot arm positioned under mesh occasionally delivered (Fig. 1a). We recorded neuronal activities using electrodes that covered the sensorimotor cortex over both hemispheres (Fig. 1b). Two cameras videotaped the behavior of the rats for simultaneous 3D tracking. Recording sessions (*n* = 106 in total) were distributed over three months and varied between 30 and 60 min (*µ* = 36.06 min, *σ* = 5.23 min). In total, we identified 3,723 single-units (*µ* = 35.12, *σ* = 20.71 across sessions) that we used for further analysis: 730, 896, and 230 in the left M2, M1, and S1, respectively, and 432, 793, and 642 in the right M2, M1, and S1, respectively (*24*).

**Figure 1:**
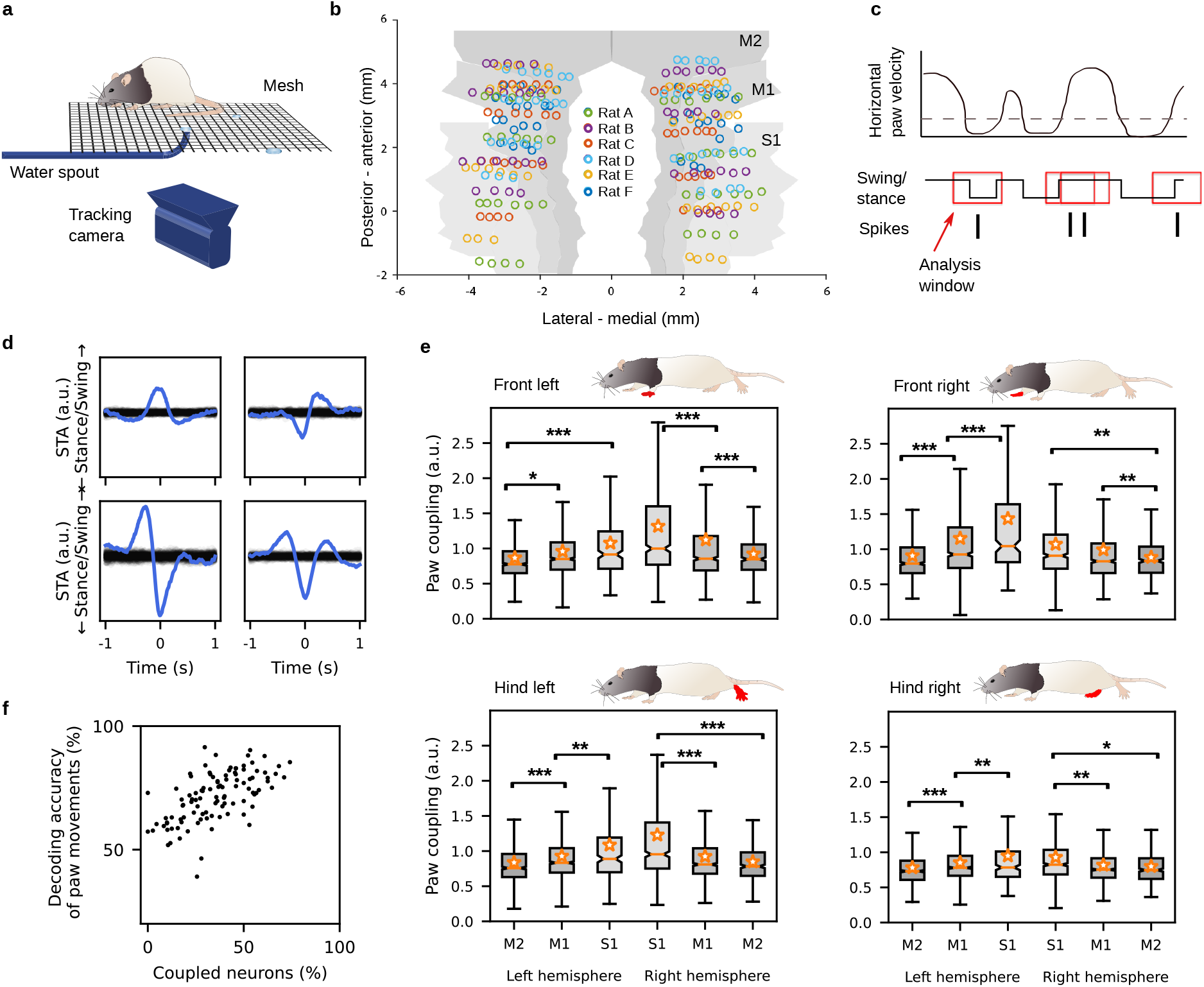
Spike-triggered average paw swing–stance status (STAPSSS) during unconstrained movements extracts lateralized paw coupling. (a) Behavioral setup with a ground mesh, camera, and robot arm delivering water drops, adapted from (*49*). (b) Locations of the electrodes of the six implanted rats, adapted from (*49*). (c) Paw movements were binarized into swing (moving) and stance (not moving). STAPSSS was calculated by averaging the swing– stance status in windows *±* 1s (indicated with red boxes) around each spike. (d) STAPSSS for the right front paw of four example single-units in the left and right S1 (upper panel) and the left and right M1 (lower panel). Black lines refer to the statistical control waveforms. (e) Coupling for each paw, brain area, and hemisphere, averaged over neurons. Stars denote the results of the post-hoc Tukey–Kramer tests (only intra-hemispheric results are indicated). Orange stars denote mean values, and notches represent the 95% confidence intervals for the median. See the main text for a definition of paw coupling. \**p* < .05, \*\**p* < .01, \*\*\**p* < .001. (f) The accuracies of neural networks trained to predict the status of the right front paw from the neural data were strongly correlated to the percentage of significantly coupled neurons.

We focused on step-like behavior to extract behavioral components from the movements. To extract the steps, we binarized the movements of the paws into swing (moving) and stance (not moving) according to a horizontal velocity threshold (0.03 mm/ms). With each paw, rats performed one step per second on average (*µ* = 1.22, *σ* = .29). The average percentage of time spent in the stance phase across rats was 71.36%, *σ* = 16.61%.

### The strongest paw coupling in contralateral S1

Since classical methods such as peristimulus time histograms (PSTHs) are not applicable to behavior without a trial structure, we computed spike-triggered averages to investigate the relationship between neuronal activity and unconstrained movements (*25*). We defined the spike-triggered average paw swing–stance status (STAPSSS) as a rough measure of the coupling of individual neurons to paw movements. For each neuron and each paw, we calculated the STAPSSS by averaging the swing–stance status in the period *±*1s around the spikes (Fig. 1c). For statistical control, we randomly shifted the spike train 1,000 times to calculate 1,000 control STAPSSS waveforms. Bootstrapping via temporal shifting preserves autocorrelations in time series and, therefore, helps to exclude false positives arising solely from autocorrelations in paw movements. We considered the STAPSSSs to be significant if their standard deviation over time exceeded the .99 quantile standard deviation of the control STAPSSS waveforms. Only neurons that spiked more systematically than expected by chance in relation to movement parameters could pass this test. Significantly coupled neurons were characterized by clear peaks in the STAPSSS (Fig. 1d, Fig. S1). In total, 54% (2,029/3,723) of all neurons were significantly coupled to at least one paw. These were 45% (534/1,162) of all neurons in M2, compared to 53% (908/1,689) in M1 and 67% (587/872) in S1.

To take into account the strength of coupling, we defined a continuous measure for paw coupling as the ratio of the STAPSSS’s standard deviation and the control standard deviation (*>* 1 for significant neurons). Using this quotient as a dependent variable, we calculated three-way ANOVAs (with hemisphere, area, and rat as factors) for all four paws separately (detailed results in Table S2). In summary, for all four paws, we found a stronger coupling on the contralateral side (*p* = .04), which suggests lateralization during locomotion. The coupling increased from anterior to posterior areas (*p <* 1e *−* 11). For all four paws, the highest mean coupling was localized in contralateral S1 (Fig. 1e). In three out of the four paws, the interaction between the area and hemisphere was also significant, that is, the differences between the contralateral and ipsilateral hemisphere increased from anterior to posterior areas (*p* = .02). To further investigate the difference in magnitude between contralateral and ipsilateral paw coupling, we defined contralateral bias as the ratio between the coupling of the contralateral and ipsilateral paws: *b* = *c_r_/c_l_* for left-hemispheric neurons and *b* = *c_l_/c_r_* for right-hemispheric neurons (*b ≈* 1 for non-biased neurons), with bias denoted as *b*, coupling as *c*, the right paw as *r*, and the left paw as *l*. We calculated this bias separately for the front and hind paws. A two-way ANOVA on the contralateral bias of individual neurons revealed a significant effect of the brain area for the front paws (*F*_2_,_3715_ = 44.66, *p <* 1e*−*19) and the hind paws (*F*_2_,_3715_ = 54.56, *p <* 1e*−*23). This confirmed that single neurons had a larger contralateral bias from anterior to posterior areas for the front and hind paws (Fig. S2).

To further investigate the temporal relationship between neuronal activity in different motor areas and paw movements, we quantified the offset between each movement peak (the STAPSSS peak) and spike. We found that for all four paws, the offset for neurons in S1 tended to be more negative (i.e., the spike followed movement) than that of neurons in M1 and M2 (Fig. S3). This effect was more pronounced for the hind paws according to an unpaired two-tailed t-test between offsets from S1 and M1/M2 (front left paw *t*_3721_ = *−*2.68, *p* = .007; front right paw *t*_3721_ = *−*2.63, *p* = .008; hind left paw *t*_3721_ = *−*6.57, *p <* 1*e −* 10; hind right paw *t*_3721_ = *−*5.94, *p <* 1*e −* 08). The finding that neurons in S1 tended to spike after movements, whereas neurons in M1 and M2 spiked in a closer temporal relationship to paw movements, aligns well with the idea of S1 reacting to sensory input and M1 and M2 being more involved in movement generation.

### Single-unit activity allows for the decoding of paw movements within sessions

Due to the strong paw coupling, we hypothesized that it is possible to decode the paw movements of freely moving rats from neuronal activity. To test this hypothesis, we applied feed-forward neural networks to decode the swing–stance status of the right front paw posed as a two-class classification problem. For each time point, we fed in the spike trains *±*400 ms of all units in time bins of 10 ms. Deep neural networks were trained and evaluated separately for each recording session. We chose this approach because single-neuron activity does not generalize over sessions, in contrast to our population-level decoding approach in the following section. The mean per-class decoding accuracies were well above chance level (*µ* = 71.47 %, *σ* = 9.98 %; chance level 50 %). While there was no significant correlation between accuracy and train set sizes (Spearman’s *ρ* = .17, *p* = 0.07), we found a significant correlation between the accuracy and percentage of coupled neurons according to our STAPSSS analysis per session (Spearman’s *ρ* = .63, *p <* 1e*−*12, Fig. 1f). This confirms that STAPSSS is a reliable measure of the correlation between neuronal activity and movement.

### The structure of population activities allows the decoding of behavior

Due to the promising decoding results of paw movements, we sought to determine whether population activities during unconstrained movements also contained information on more complex behavior. We used LEMs (*22, 5*) to reveal and visualize the structures in the population activities. LEM is a non-linear dimensionality-reduction method for extracting low-dimensional manifolds in high-dimensional data using spectral techniques. We applied LEM to the neighborhood graphs of neuronal activity vectors to visualize structures and relationships among population activities at different time points of a recording session in a low-dimensional space. Most of the resulting projections showed a clear saddle-like shape when visualized in three dimensions (Fig. 2a, 52/95, roughly 55% of session structures had a similar shape, as classified visually), although there were also random-like population structures that differed from the majority (14/95, about 15% of the sessions, Fig. S4; the remaining 35% had intermediate levels of structuredness). In random-like population structures, the time points were uniformly distributed in a sphere.

**Figure 2:**
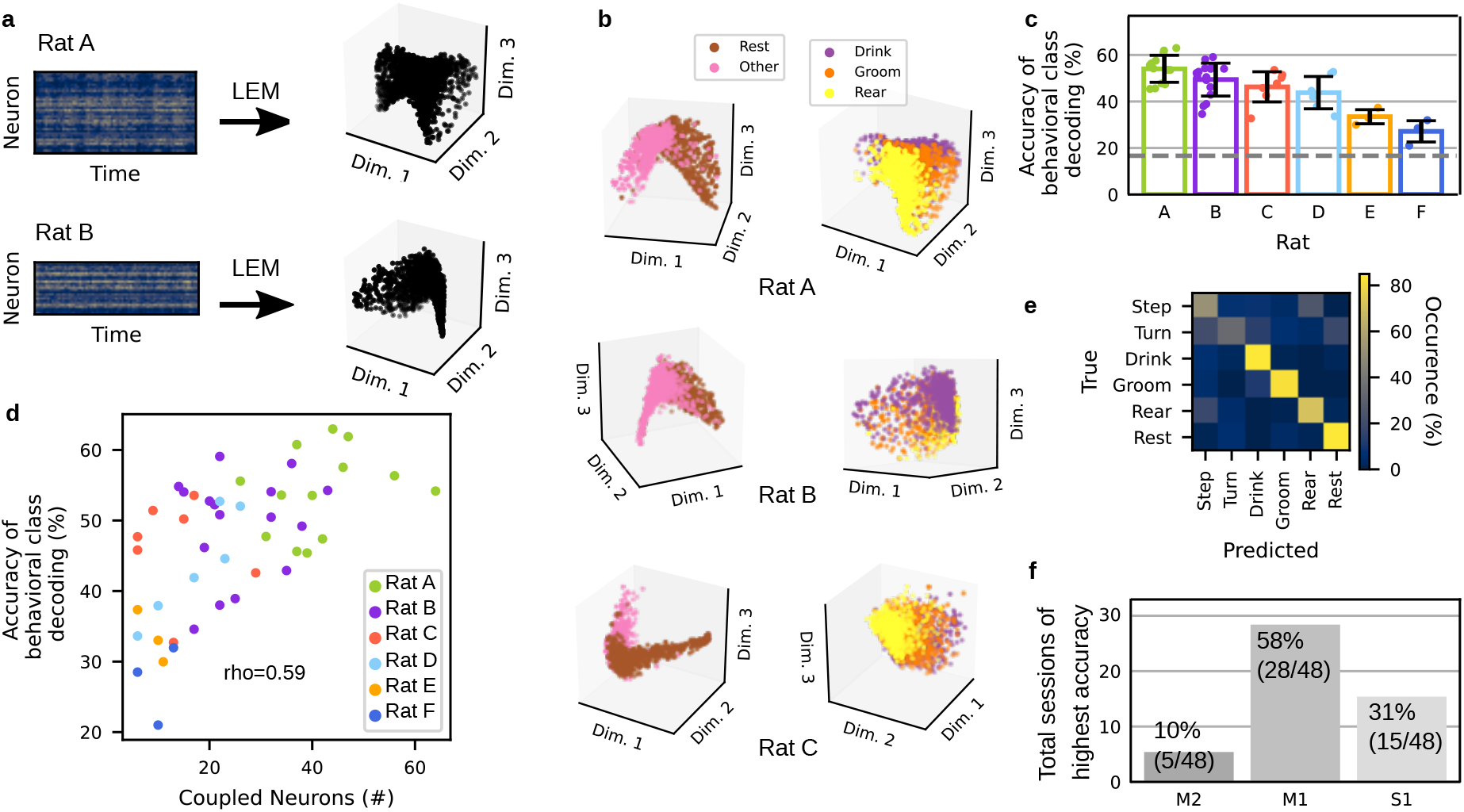
Behavioral classes can be decoded from low-dimensional neural structures within one session. (a) Non-linear dimensionality reduction through LEM was performed on the neural data of each session separately. (b) In the low-dimensional space, different behaviors were distinguishable in as few as three neural dimensions. Left panel: The first dimension clearly differentiated between rest and movement (all other behavioral classes). Right panel: The second and third dimensions played a primary role in coding the difference between paw- and head-related behavior (rear vs. drink). One session for each of Rats A, B, and C is depicted. (c) Classification accuracies for the six behavioral classes given low-dimensional neural input were above chance level for the sessions for all six rats. The gray dashed line indicates the chance level, and the error bars show standard deviations. (d) Accuracies were correlated to the number of significantly coupled neurons (neurons coupled to at least one paw according to the STAPSSS measure). (e) One example confusion matrix for the test set of a single session of Rat A, with a mean per-class accuracy of 68.46%. (f) For most of the sessions, classification accuracies for the six behavioral classes were the highest given dimensionality-reduced neural activity from M1 as input, followed by S1 and M2.

Sessions with a clear saddle-like shape were characterized by a larger number of neurons that were significantly coupled to at least one paw compared to sessions with an intermediate or low level of structuredness (23.57 *±* 14.71 vs. 16.62 *±* 12.33 neurons, *t*_93_ = 2.43, *p* = .016). To ensure that the saddle-like structures were not a simple artifact of the dimensionality-reduction method, we also performed time-shuffled, neuron-shuffled, and time-shifted control reductions (*5*). These did not lead to any apparent structure (Fig. S5).

To investigate the relationship between population structures and the corresponding behavior, we proceeded by manually labeling sessions in 500-ms snippets into six behavioral classes (stepping/paw movement, turning/head movement, drinking, grooming, rearing, and resting). We included all sessions with clear saddle-like shapes and with at least five significantly coupled neurons, which resulted in a total of 48 sessions (13 for Rat A, 16 for Rat B, 7 for Rat C, 6 for Rat D, 3 for Rat E, 3 for Rat F).

While each session contained at least some samples of each behavior, the occurrences of behaviors still differed considerably across sessions and rats (Fig. S6). In contrast, the distributions of behaviors across the neural structures revealed clear similarities across rats, which was surprising assuming a sampling of approximately .005% of all neurons^1^ on average in only roughly overlapping recording sites (Fig. 1b). For example, the second eigenvector (here: first dimension), the so-called Fiedler vector, clearly represented the difference between movement and rest (Fig. 2b left column). For some animals, a clear distinction between more paw-related (paw movement, rearing) and head-related behavior (head movement, drinking) was observable in the third and fourth eigenvector (here: second/third dimension, Fig. 2b right column). Although the position of a population vector in the LEM space is univocally defined by the instantaneous activity of all its composing units, and is relatively little affected by the activity of a single-unit, there is a relationship between the overall structure emerging in the LEM space when observing the totality of recorded data and the firing of neurons with high behavioural selectivity. While population vectors cluster in space due to the similarity between neuronal representations during a specific behavior, single-units with high selectivity for such a behavior will fire more strongly at that behavior’s cluster (Fig. S7). Moreover, the distance in the LEM space between population vectors corresponding to two behaviors will increase with the number of units within the population that change their firing rate between the two behaviors (Fig. S8 a,b).

To quantify the separation between the neuronal representations of the six identified behaviors in the LEM space, we trained a neural network based on the first 10 dimensions of the population vectors. We chose 10 dimensions because we found the mean dimensionality in the LEM space to be 8.59, *σ* = 1.11 (see Methods). By choosing a slightly higher value than the mean dimensionality, we added a small safety margin to ensure the inclusion of all relevant dimensions. The neural network correctly classified behaviors more frequently than by chance (mean per-class accuracy *µ* = 47.11%, *σ* = 9.62%; chance level 16.66%, Fig. 2c). The accuracies were correlated to the number of significantly coupled neurons (*n* = 48, Pearson’s *ρ* = .59, *p <* 1e*−*5, Fig. 2d), the total number of units (Pearson’s *ρ* = .54, *p <* 1e*−*4, Fig. S9a), and the signal-to-noise ratio (SNR) averaged over units (Pearson’s *ρ* = .49, *p* < .001, Fig. S9b). Common classification mistakes consisted of confusing rearing or turning with stepping, as well as turning with drinking or resting (Fig. 2e). At the single-unit level, these behavioral classes shared, in fact, the highest number of selective units (Fig. S8 c,d). Moreover, we observed the lowest accuracy for Rat D, Rat E, and Rat F. These rats had a low mean SNR (Rats E-F, Fig. S9b) or no electrode coverage of posterior areas (Rats D and F, Fig. 1b). This last aspect made us hypothesize that more posterior regions are primarily involved in the encoding of behavioral classes. To test this hypothesis, we investigated the influence of the different sensorimotor areas on neural population structures. Thus, we conducted dimensionality reductions with equal numbers of neurons (i.e., 20 randomly chosen units) from M2, M1, or S1 as input. With this subset, we trained artificial neural networks to decode the behavioral classes with the neural activity in a given area reduced to five dimensions as input. The decoding accuracies for M1 were significantly better than those for M2 (paired two-tailed *t*-test, *t*_40_ = 4.18, *p* < .001) and slightly, but not significantly, better than those for S1 (*t*_41_ = 1.90, *p* = .06). In total, the accuracies were highest in M1 for 28 out of the 48 sessions, compared to 15 for S1 and 5 for M2 (Fig. 2f, accuracies *µ* = 25.80 *±* 4.92% in M2, *µ* = 28.60 *±* 5.34% in M1, *µ* = 26.66 *±* 5.41% in S1). The low relevance of anterior sensorimotor regions is in line with the STAPSSS results, as well as with the lower decoding accuracies in Rat D and Rat F.

### A cross-session polytope comparison reveals similarities in the average encoding of behavioral classes across animal

The visual similarity between neural population structures in three dimensions (cf Fig. 2a,b) led us to wonder whether correspondences between the full dimensional structures could be quantified. Such similarities become more apparent when reducing the extended manifolds to polytopes – high-dimensional polyhedra – with vertexes defined by the average population vectors associated with the six behavioral classes (Fig. 3a). This encouraged us to systematically investigate whether population activities during unconstrained movements contained structures that were conserved across recording sessions or even across different animals. We excluded Rat F from all of the following analyses because of low recording quality, which may reflect the long delay between implantation and measurements compared to the other rats (see Table S1).

**Figure 3:**
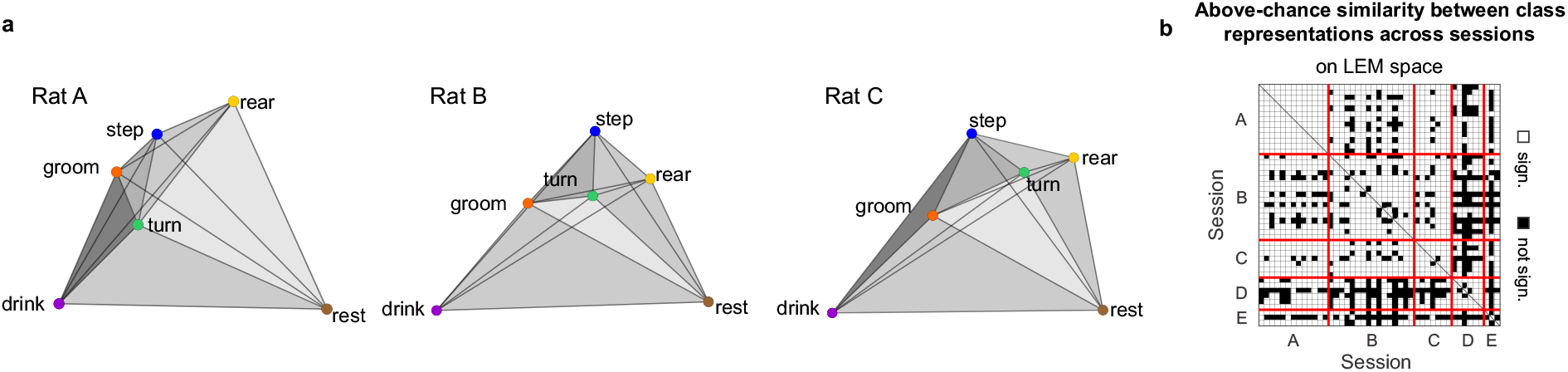
Cross-session polytope comparison reveals similarities in the average encoding of behavioural classes across animals. (a) 3D polytopes in the LEM space identified by the average population vectors of the six behavioral classes for one example session of Rats A, B, and C, respectively. The distances between the polytope vertexes are proportional to the distances computed between the average population vectors in the 3D LEM space. The gray shading is added to visualize the 3D structure. (b) Significant similarities among the polytope structures of different sessions. Similarity was tested for each pair of sessions by comparing, across-sessions, the difference between the session-specific matrices collecting the Euclidean distance between the average population vectors associated with each behavioural class (see Methods). Significance was assessed by bootstrapping the class labels (n = 720, all possible permutations of class labels). Distances were computed in the 20-dimensional LEM space. Of the 990 possible session pairs, 78% had a p-value below 0.05, indicating that the similarities of the neuronal activities could be captured using the polytope structures.

Polytopes are useful tools to facilitate the visualization of the complex structure associated with the neuronal representations of the six identified behaviors and their reciprocal distance. To test whether such distances were preserved across sessions and animals above chance level, we first tested whether behaviors associated with similar population vectors in one session corresponded to behaviors with similar population vectors in other sessions. For example, if in one session the population vectors during turn and step are similar to each other but dissimilar from those during rest, we wondered whether the same relationship can be found in other sessions as well. More formally, for each pair of sessions *v* and *w* and each behavioral class *i*, we ranked the remaining classes by the Euclidean distance between their average population vector and the average population vector of class *i*. If *v* and *w* had the same polytope structure, the rank associated with each behavior would be identical. We quantified the similarity between ranks across sessions with the statistic *s^vw^*, defined as the number of concordant ranks, and compared its distribution with that obtained from bootstrapping (see Methods for details). We found significant similarities across sessions, both when computing distances in the high-dimensional recording space (Kolmogorov–Smirnov test, *p <* 1*e−* 39) and in the reduced LEM space (*p <* 1*e −* 73) (Fig. S10 a-b). This finding confirms that the obtained result was not an artifact of the dimensionality-reduction procedure. Moreover, to ensure that such significance did not depend exclusively on the enhanced distance between the “rest” class and any other classes, we repeated the analysis with “rest” excluded from the accounted classes (*p <* 1*e −* 21, Fig. S10c).

We performed a second test to compare the overall conservation of relative distances among the population vectors associated with different behaviors at the single-session level. We captured the differences between the cross-behavioral distance matrices of two sessions with the Jeffries–Matusita metric and compared them with the bootstrap distribution obtained by shuffling the behavioral labels (see Methods for details). This was done for each pair of sessions within and across animals, both in the high-dimensional recording space and in the LEM space, and by excluding the “rest” class from the test (Figs. S10d, 3b, and S10e, respectively). In all of these cases, the similarity between the polytopes of different sessions and animals was above chance for most session pairs, and non-significance often occurred for animals with a low SNR (cf. Fig. S9b).

### Cross-subject and cross-session decoding

Polytopes capture the distance between the average neuronal representations of the different behaviors but neglect their shape and extension on the manifold. Encouraged by the similarities observed among the polytope structures of different recording sessions, we decided to perform a stronger test and attempted cross-subject decoding. Cross-subject decoding requires not only an agreement between the average representation of behaviors but also accounts for the variability in neuronal representations associated with each behavior. While it is impossible to find a direct correspondence at the single-neuron level across animals, similarities in lower-dimensional population structures can be used for cross-subject and cross-session decoding (Fig. 4a). For the decoding analysis, we divided the six behavioral classes (Fig. 4b) into two disjointed sets: one “alignment set”, which was used to align the neural structures, and one “decoding set”, which was used for training and testing a classifier. Thereby we ensured that no class was used for aligning the structures and classification in the neural space at the same time. The mean neural vectors (four dimensions) corresponding to the behavioral classes in the alignment set were used to compute a Procrustes transformation between two sessions to align the population activity structures (*27, 28*) (Fig. 4c). Procrustes transformations involve translation, scaling, reflection, and rotation and thus preserve the shape of a set of points. For decoding, we trained a classifier on samples from the decoding set of one session for a single rat using the activity in the dimensionality-reduced neural space as input. Then, we tested the generalization on another session of the same rat (cross-session decoding) or another rat (cross-subject decoding) (Fig. 4c). Notably, the samples of the decoding set of the two tested sessions were not used for computing the Procrustes transformation. In the first experiment, the alignment set consisted of four behavioral classes, with two other classes remaining for the decoding set. This resulted in a total of 15 possible splits into two sets. Classifiers trained on highly decodable sessions also successfully generalized to other sessions from the same or other rats (Fig. 5a–c, Fig. S13a–b). In the generalization matrix (Fig. 5a), 13.88% of the generalization results (275 out of 45*44=1980) had a mean per-class accuracy higher than 60%, and 59 higher than 65%. In the set of sessions with the highest 10% signal-to-noise ratio (SNR *>* 4.33), the mean per-class accuracy in the generalization task was *µ* = 59.72%, *σ* = 5.37. The best-performing sessions included sessions of Rats A, B, and C with sufficient recording quality and a sufficiently high number of units for a robust estimation of the underlying population structures (Fig. 5d). Additionally, the correlation between within-session and between-session accuracies was high (Fig. 5c, *n* = 45, Pearson’s *ρ* = .68, *p <* 1e*−*6). We defined the “generalization accuracy” of a session as the average test accuracy across all sessions (mean value per row of Fig. 5a first matrix). These generalization accuracies were correlated to the total number of units (Pearson’s *ρ* = .38, *p* < .01), with a higher number of units leading to a better estimation of the population structure. The generalization accuracies were also correlated to the session length (Pearson’s *ρ* = .37, *p* < .05) since the number of samples used for LEM (which included only time points with sufficient activity) varied across sessions and rats. Finally, the recording quality—namely, the SNR averaged over units—was correlated with generalization (Pearson’s *ρ* = .38, *p* < .01). Particularly, Rats A and B, which performed best in the generalization, had both a high SNR and a high total number of units (Fig. 5d).

**Figure 4:**
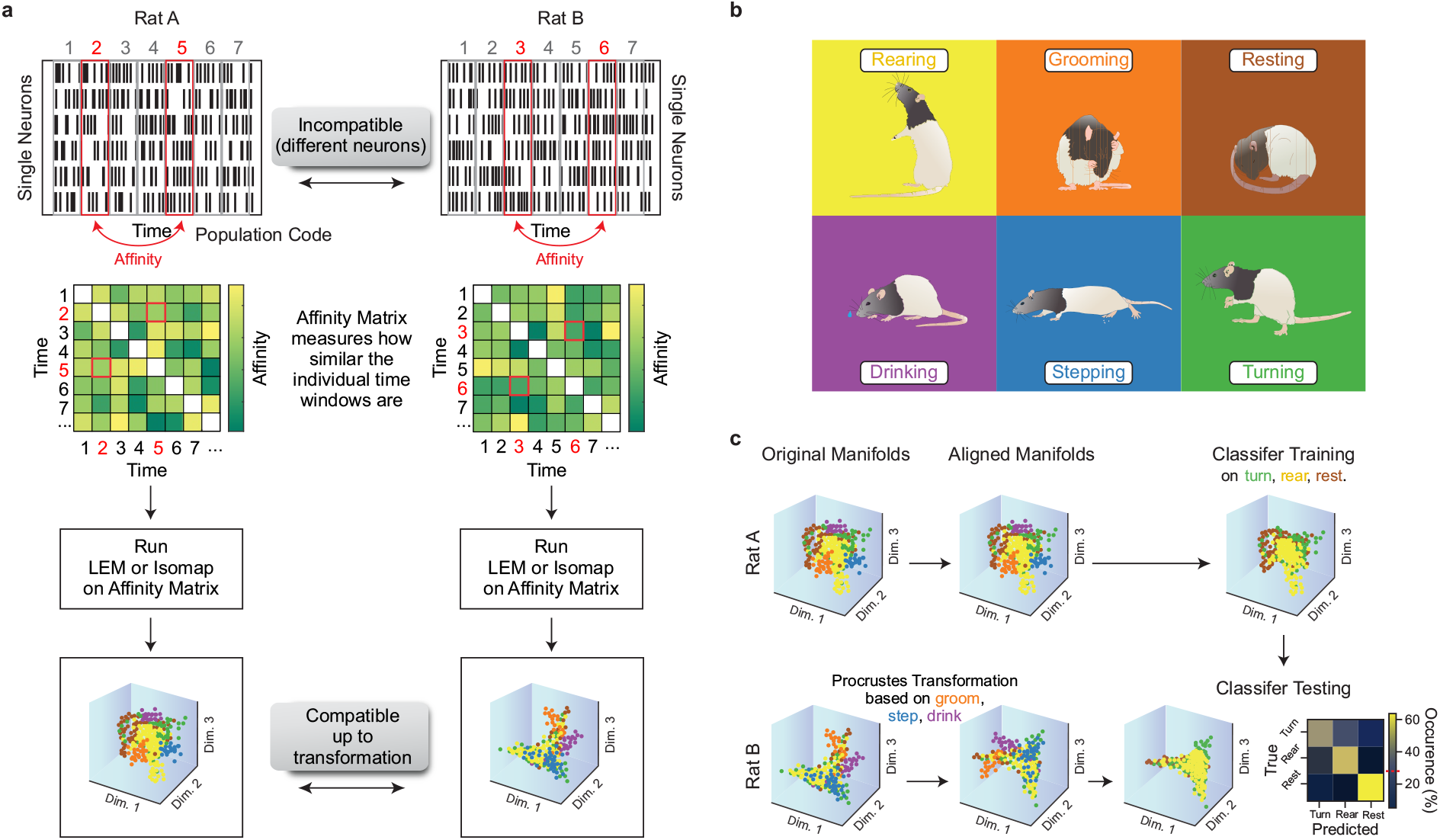
Alignment procedure of neural structures for cross-subject generalization. (a) Neuronal activity on the single-neuron level is not comparable across rats and sessions. To reveal population structures that are compatible—up to a transformation—across subjects, we computed the affinities of the population activity at different time points. The resulting affinity matrices could then be used for dimensionality reduction. (b) This represents the different classes of behavior exhibited by the rats. (c) We aligned the low-dimensional neuronal structures of two different sessions using a Procrustes transformation. This procedure used only a subset of the behaviors (e.g., groom, step, and drink). After alignment, a classifier trained on one rat could generalize to another. In the classification, another subset of behaviors (e.g., turn, rear, and rest) was used.

**Figure 5:**
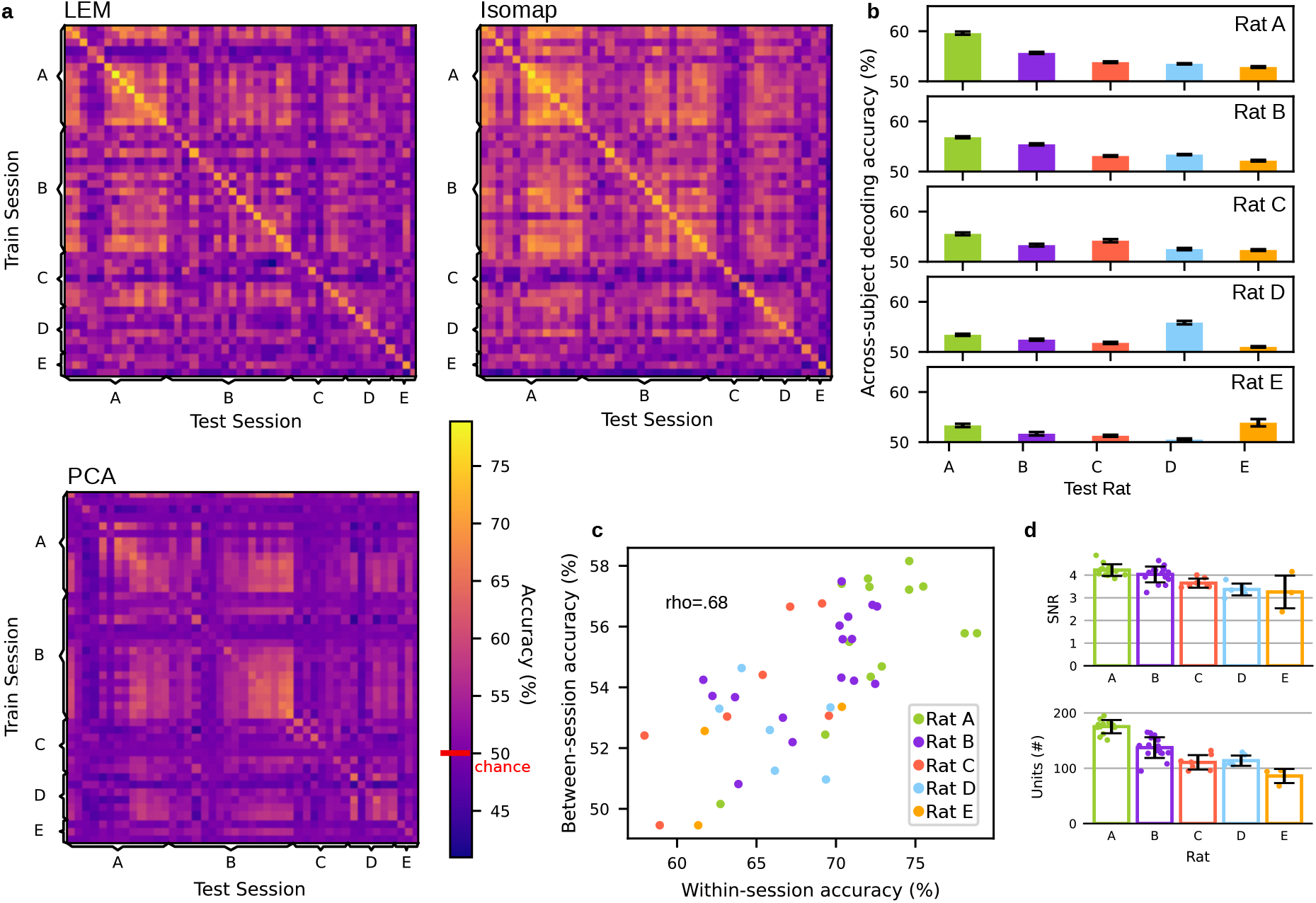
Similar structures of population activities allow for cross-subject decoding. Results for a classifier trained on two behavioral classes of one rat (chance level 50%, red line) and tested on another rat after using four disjointed behavioral classes to align the neural manifolds. All values in the orange-to-yellow spectrum indicate accuracies above the chance level. (a) Complete mean per-class accuracies across training and test sessions when aligning on four classes and testing on two in the LEM/Isomap/PCA spaces. On the diagonal, training and test data came from the same session. Off-diagonal entries refer to tests on data sets that were not identical to those of the training session. Values are averaged over 20 runs, and there are 15 possible splits of the six behavioral classes into alignment and decoding sets. (b) Average of the mean per-class accuracies when training a classifier on the rat indicated by the row and testing on the rat indicated by the column. Error bars provide the standard error of the mean. (c) Within-session and between-session accuracies were highly correlated; sessions that were easier to classify generalized better to other sessions. (d) The best-performing rats (Rats A and B) were also those with the highest number of recorded units and the highest SNR. Conserved structures of neural activity in sensorimotor cortex of freely moving rats allow cross-subject decoding –

### Decoding is robust to methodological and class-selection changes

To evaluate whether our results depended on a specific dimensionality-reduction method, we used Isomap, another non-linear dimensionality-reduction method. Also Isomap revealed neuronal structures that were comparable across subjects (Fig. S11, Fig. 5a second matrix). In contrast, linear methods such as principal component analysis (PCA) were not powerful enough to extract these neuronal structures (Fig. 5a third matrix); the LEM results were significantly better than those from data reduced using PCA (*t*_30374_ = 40.33, *p* = 0).

For a more systematic test of the relationship between the number of units and generalization, we took all sessions with a generalization accuracy of at least 55% (a total of 19 sessions from Rats A, B, and C) and conducted an ablation study with LEM reductions on the reduced number of units (20, 40, 60, and 80 units removed per session). We then repeated the generalization experiment on the aligned LEM structures. The accuracies steadily decreased with fewer units (Fig. S12), confirming the high relevance of the number of units for a robust estimate of the population structure.

To determine which sensorimotor areas were most relevant for generalization, we again took the 19 best sessions and conducted LEM reductions after removing M1, S1, or M2. Additionally, for a fair comparison, we removed a random portion of all of the other units for underrepresented areas such that the number of units after the removal of M1, M2, and S1 neurons remained constant across sessions. Generalization accuracies on aligned LEM structures decreased considerably after the removal of M1 (accuracies *µ* = 53.91 *±* 7.48%); these values were slightly but significantly lower than after the removal of units from S1 (accuracies *µ* = 55.94 *±* 7.73%, paired two-tailed *t*-test, *t*_5414_ = *−*16.07, *p <* 1e*−*56), M2 (accuracies *µ* = 54.99 *±* 8.32%, *t*_5414_ = *−*8.14, *p <* 1e*−*15), or of the same number of units distributed over all areas (accuracies *µ* = 55.55 *±* 7.84%, *t*_5414_ = *−*13.89, *p <* 1e*−*42).

In a second experiment, we used only three classes in the alignment set and the three remaining in the decoding set to test the generalization under more difficult conditions, resulting in 20 possible splits of the six classes in total. The general pattern of the generalization matrix stayed the same (Fig. S13c–d). To verify that the classifiers did not only learn to discriminate the simplest difference—the difference between rest and movement—we conducted another experiment without the class “rest”. Although the accuracies were lower in this setting, the general pattern remained the same (Fig. S13e-f). To assess the relevance of the alignment of neural structures, we also tested the generalization on neural structures without explicit alignment as a control. In most cases, the accuracies on aligned structures were much higher than those on unaligned structures (Fig. S14).

To further explore our results, we performed control classification experiments in which we examined the generalization on shuffled data (see Fig. S5). Accuracies of shuffled data were significantly lower than those computed in the original LEM space (*t*_30374_ = 82.31, *p* = 0 when comparing with neuron-shuffled, *t*_30374_ = 83.32, *p* = 0 with time-shuffled, and *t*_30374_ = 75.35, *p* = 0 with time-shifted data) and did not exceed the chance level (Fig. S15). Furthermore, we computed LEM reductions in the non-binarized neural space and repeated the generalization experiment. Also in this case we could find significant generalization for multiple sessions, but with accuracies lower than when the analyses were performed on binarized spikes (Fig. S15d, *t*_30374_ = *−*28.72, *p <* 1e*−*178). This supported our intuition that an analysis seeking to identify similarities in the neural space between sessions should not be biased toward more active neurons.

## Discussion

In this study, we investigated single-neuron activity as well as population activity patterns in the rats’ sensorimotor cortex during unconstrained and self-paced behavior. The behavior was as closely related as possible to naturally occurring behavior, as it was based on foraging, but it was still performed in a limited arena to allow for reliable movement tracking. The first analyses were sanity checks to validate our approach of studying freely moving animals without a clear trial structure. Based on the chosen measure, STAPSSS, 54 % of all neurons were significantly coupled to paw movements. This fraction of coupled neurons is in the range of previously reported numbers. For example, 60 % of neurons in the hindlimb motor cortex reacted to different locomotion scenarios (*29*), and 44 % in M1 were body-coupled in freely moving rats (*25*). Our multi-side recording approach allowed us to comprehensively test for differences in neuronal activity across the entire sensorimotor cortex. Previous research has found that the laterality of forelimb representations increases from M2 to M1 in a pedal task for head-restrained rats (*30*). Here, we extended this laterality gradient to more posterior regions— in particular, S1. As we targeted the output layer of the cortex (layer V), we putatively biased our recordings toward pyramidal tract neurons, which have been described as being predominately involved in laterality (*30*).

While the above-described findings refer to the general features of the sensorimotor cortex, the main finding of our study was based on conserved neuronal population structures. Experimental, computational, and theoretical work has identified a rich structure within the coordinated activity of interconnected neural populations in movement control, decision-making, and memory tasks. These findings are conceptualized within the framework of neural population dynamics, which can reveal general motifs (*2*). Recurrent neural networks (RNNs) can be applied to neural data to reveal structural and geometric properties (*31*). Multiple tasks can then be represented in different RNN models. In these networks, some clusters of units have been identified as specialized for subsets of tasks (*1*). Alternatively, methods such as PCA and its variants dPCA and jPCA have been applied to identify the stability of motifs across modalities such as arm and speech control (*3*), as well as within and across brain areas (*4*).

In contrast to the previously described studies, we focused on the existence of conserved neuronal structures across animals without any clear instructed task line but with several behavioral classes. These two points differentiate our study from previous publications in the field. We investigated population activity patterns, which are commonly assumed to reside on low-dimensional manifolds in the full neural state space (*14, 16, 32, 33, 34*). In contrast to the (globally) linear method like PCA that most studies have used (*12, 20, 15, 16, 4, 17, 18, 35*), we assumed the preservation of local neighborhood relations in the data. Therefore, we employed LEM (*22, 5*) to reveal the presumed preserved low-dimensional structures. Remarkably, neuronal population activity during unconstrained behavior contained similar structures across animals and sessions, already visible in the first three dimensions. Furthermore, the distribution of different behaviors across low-dimensional neural structures was systematic, which we confirmed with our above-chance, within-subject decoding results. The allocation of different behaviors on the population structures revealed strong similarities across rats. Particularly, movement and rest could be clearly visually distinguished in the first dimension. This is in line with results on clear separations in the neural state space for output-potent and output-null (e.g., preparatory) neural activity (*12, 20*).

To support our main claim that low-dimensional neural manifolds are comparable across sessions and animals even in the case of unconstrained behavior, we first showed with our polytope analysis that the relative positions of the neural representations associated with different behavioral classes were conserved across animals and sessions above chance. The analyses of the polytope structures compared the distance between the average neuronal representations of behaviors, neglecting their precise spatial extension on the manifold as determined by the variability in neuronal representation of each behaviour. Conversely, cross-subject decoding was also affected by such variability and, therefore, tested an even stronger degree of similarity. Ultimately, we evaluated the performance of a classifier trained on the neuronal activity of one subject to predict the behavior of another based on its own neuronal activity; this provided us with a proxy to experimentally test and quantify the degree of universality of mental representations across subjects. Since the neuronal state space of different subjects cannot be directly compared (given the difference in number and identity of the recorded neurons), applying dimensionality reduction and alignment was necessary to achieve this goal. A simple, supervised, shape-preserving alignment procedure—namely, a Procrustes transformation between mean population vectors for different behavioral classes in the dimensionality-reduced neural space—sufficed for successful cross-subject generalization in a decoding task with distinct but related behavioral classes. Our procedure was applicable to sessions with sufficient recording quality (indicated by a high SNR of the recorded units) and enough units for robust population estimation. Further, the generalization accuracies of the sessions were closely related to their within-session accuracies. Generalization was considerably worse for population structure estimates based on fewer units. In line with the within-session decoding results, we also found that generalization significantly decreased after the removal of M1, which indicated consistent population responses especially in this area. The low relevance of the anterior motor cortex to information regarding behavioral categories is in line with our STAPSSS results. Nevertheless, in contrast to the encoding of paw movements, our results on the population decoding of the higher-level behavioral categories hinted at major contributions from M1, not only S1. Thus, our results close a gap in a previous study that investigated postural and behavioral encoding in the posterior parietal cortex and M2 (*36*). While we mostly used LEM as a dimensionality-reduction method given its solid theoretical basis, we also showed that another non-linear dimensionality-reduction method, Isomap, can be used to reveal neural structures that are comparable across subjects.

A shared structure in neuronal activity across subjects has been shown mostly in fMRI studies, where even between-subject classification has been demonstrated (*37, 27, 38, 39, 40, 41*). While these works have focused on watching movies (an activity that can be conducted similarly for different subjects), EEG and EMG cross-subject decoding has been shown for hand movements (*42, 43*). In rodents, related work has shown the cross-subject decoding of odor sequences in the orbitofrontal cortex (*44*) and place-cell activity in the hippocampus (*45*). In contrast to these studies, we showed cross-subject classification in a more complex case where rats roamed freely without training or a trial structure in the underlying task. Therefore, our main finding of shared neural structures is consistent with recent findings but also extends them to more complex, less constrained behavior. To our knowledge, this is the first time that the conservation of neural structures across animals and for distinct, spontaneous behavioral classes has been shown. This finding implies that conserved neuronal structures occur without training. Therefore, the neuronal computations underlying these structures might be similarly realized across individuals, either from birth or during development.

Remarkably, sampling as few as approximately 0.005% of all neurons in only roughly overlapping electrode positions sufficed to estimate population structures that were similar enough to allow for cross-subject generalization, at least for sessions with a sufficient number of units to allow for robust neural manifold estimation. Internal states (such as thirst, attention, or motivation), which we did not analyze here, may also have influenced the neuronal activities (*46*). It has been hypothesized that cross-individual decoding might not be possible with increasing task complexity (*5*). However, our results indicate that even during unconstrained behavior, the relationships among neural activity patterns are conserved across different animals. This conservation of population-level neural phenomena provides a foundation for cross-subject decoding, even in the difficult case of unconstrained behavior.

## Acknowledgments

We thank Krishna Shenoy, Philipp Schröppel, and Christian Zimmermann for their comments on the manuscript.

## Author contributions

T.B. and I.D. designed the study. S.M. analyzed the data, except the parts that E.R. analyzed. E.R. performed the polytope and behavioral single-unit coding analyses. D.D. contributed to the design of the analyses and the statistical evaluation. S.M., T.B., I.D, E.R., D.D. and A.S. wrote and edited the manuscript. D.E. conceived and recorded the data set.

## Data and materials availability

Code for the most important functions is available at https://github.com/Optophys/Conserved_structures_cortex. Data is available from the authors upon reasonable request.

## Supplementary materials

Methods

Figs. S1 to S15

Table S1 to S2

Supplementary Materials

### Methods

#### Animal surgery

We implanted six male Long Evans rats at the age of eight weeks with 22 tungsten electrodes (200 to 600 kOhm impedance, polyimide insulation, WHS Sondermetalle, Grünsfeld, Germany) at a 1.2 mm implantation depth in each hemisphere (implantation: January 2017 for Rat F, April 2017 for Rats A–E). Electrode locations spanned from -2 to +5 mm in the anterior/posterior direction and from 1 to 4 mm in the lateral/medial direction. This resulted in three medial–lateral rows of six electrodes each, plus one row of four electrodes (see Fig. 1b). Details of the procedure are described elsewhere (*49*). The Regierungspräsidium Freiburg, Germany approved all animal procedures.

#### Behavioral task

The rats were kept water-restricted for the time course of the experiments (free access to water for two days per week). For the experiments, the rats moved unconstrained on a mesh of 30*×*40 cm in a closed arena. Every 10 to 30 s, a waterspout pseudo-randomly positioned by two servo motors released a drop of water onto the mesh, which the animals could find and consume. To prevent the rats from merely following the movements of the waterspout, we included dummy movements that were not followed by a release of water. Even experienced animals were not able to predict the position of water drops without an active search, and the animals did not find all water drops throughout a session. This task has been previously described (*49*). Here, we only used part of the data set discussed in (*49*); in particular, we only included sessions with a minimum duration of 30 min.

#### Data acquisition and the preprocessing of extracellular recordings

Extracellular signals were recorded at 30kHz and band-pass filtered, amplified, and digitized using a head stage (Intan Technologies, Los Angeles, California) situated at the head of the animal. Spike sorting was conducted on high-pass filtered signals (cut-off at 300 Hz) separately for each electrode. Spikes were defined as amplitude threshold crossings of four times the standard deviation of the signals. For each spike, we extracted the window of -0.5 to 2 ms around the peak amplitude (resulting in 76 values per spike). Spike sorting consisted of two phases for each unit. First, a seed spike was estimated. This was accomplished by calculating the spike neighborhoods (spikes within the average noise level, half a millisecond before the spike, across all units) for 500 randomly chosen spikes. The spike with the most neighbors was chosen as the seed spike. Second, we optimized the spike waveform through an iterative procedure. This was done by alternating the calculation of a new noise level for the neighboring spikes, the update of the neighborhood (spikes within the new noise level), and the update of the average waveform. This iterative procedure ended when the neighborhood assignments remained constant. The algorithm proceeded with the remaining spikes by choosing a new seed spike. Details of the offline denoising and spike sorting procedure have been described elsewhere (*49*). For our single-unit analysis, we only kept single-units according to the distribution of inter-spike intervals. single-units with a firing rate lower than 0.1 Hz were not included in the analysis. Two cameras (Stingray, F033C IRF CSM, Allied Vision Technologies) positioned below the mesh tracked the movements of the colored paws. The videos were taken with a frame rate of 80 Hz and smoothed with a Gaussian filter before analysis.

#### Single-unit STAPSSS analysis

Paw movements were labeled as “swing” for horizontal velocities higher than 0.3 mm per 10 ms (the bin size we used for our analysis) and “stance” otherwise. Spikes were also binned with a bin size of 10 ms. For each neuron and each paw, we defined the spike-triggered average paw swing–stance status (STAPSSS) as the behavioral average over all windows *±*1 s around the spikes. We normalized each STAPSSS waveform by the mean. We defined the paw coupling of a neuron as the ratio of the standard deviation of the STAPSSS waveform to the statistical control standard deviation. The latter was defined as the .99 quantile standard deviation of a distribution constructed out of the standard deviations of the STAPSSS waveforms of 1,000 randomly shifted spike trains. If a neuron was not related to a paw’s movement, its STAPSSS waveform would be flat and its standard deviation would not exceed the control standard deviation. We defined the contralateral bias as the ratio of contralateral to ipsilateral paw coupling. Statistical analyses were done using the *anovan*, *multcompare*, and *ttest* Matlab functions. The ANOVA tests always included the rat’s ID as an additional factor.

#### Decoding from spike trains

We used fully connected neural networks with three hidden layers of 500 units each for decoding. The networks’ inputs were the Gaussian-smoothed (*σ* = 20ms) binned spikes in *±*400 ms, resulting in 81 input bins for each neuron. In contrast to the STAPSSS analysis, where only single-units were considered, we used all units as input for decoding. Each session was split into training, validation, and test sets (70/15/15 %). Two of the 106 sessions were excluded from decoding because of insufficient data. Training was conducted with the Adam optimizer (*47*), batch size 64, and an initial learning rate of 0.0001. A dropout rate of 75 %, L2 regularization (*λ* = 1*e −* 4), and early stopping were applied to prevent overfitting. To deal with class imbalance, we used weighted cross-entropy loss to put more weight on the less frequent class (swing). The reported accuracies were mean per-class accuracies. The decoding accuracies of the deep neural network were significantly better than a baseline linear classifier (two-sided paired *t*-test, *t* = 6.55, *p <* 1e*−*8). For the baseline, we used a logistic regression with three-fold cross-validation of the L2 regularization strength on the concatenated training and validation sets. The test sets for each session were the same as for the artificial neural network. Class weights were adjusted to be inversely proportional to class frequencies, as for the artificial neural network. The artificial neural network was implemented in Tensorflow. For the linear baseline, we used Python’s scikit-learn function *LogisticRegressionCV*.

#### Dimensionality reduction

We used LEM (*22, 5*), an unsupervised non-linear dimensionality reduction method, to investigate the low-dimensional structure of population activity. For each session, spike counts were binned in 100 ms bins and then binarized (1 for at least one spike per bin, 0 for no spikes). Single and multi units were used. Only time points with at least 15 active units were retained. Since we restricted further analysis to sessions with at least 5,000 valid time points, we considered only 95 of the 106 sessions. For each session, we constructed an unweighted, mutual kNN graph based on the Hamming distance on the columns of the *n × t* matrix (*n* units, *t* time points). Our code for LEM was built on recent work (*5*). Two iterations of the LEM algorithm were performed. However, in contrast to Rubin et al., we used the Hamming distance in the first iteration and reduced to 20 dimensions. In the first iteration, we used 0.5% of the time points as neighbors; in the second, this parameter was set to 7.5%. Furthermore, we applied a random walk normalized graph Laplacian instead of the symmetric normalized graph Laplacian, as proposed in a previous study (*48*). In detail, we constructed the unnormalized graph Laplacian as *L* = *D − W*, with *D* as the diagonal degree matrix and *W* as the adjacency matrix of the kNN graph. Solving the generalized eigenvalue problem *Lv* = *λDv* corresponded to finding the first eigenvectors of the random walk normalized graph Laplacian *L_n_* = *D^−^*^1^*L* (*48*). Since the eigenvector corresponding to the smallest eigenvalue (zero) is constant, we discarded the first dimension of the LEM for all analyses and decoding studies. The other LEM eigenvectors (=dimensions) were ordered by eigenvalue magnitude— that is, the “splitability” of the time points in different clusters (i.e., the dimensions that best divided the time points into clusters came first.) For the LEM reductions on units from different sensorimotor areas, we randomly chose 20 units from each area as input (if fewer than 20 units for an area were available, the analysis was omitted). We chose to reduce to six dimensions in the LEM space, leaving us with five dimensions for decoding with deep neural networks (as mentioned above, the first dimension of the LEM must be discarded). For the ablation study on sessions with 20, 40, 60, or 80 units removed, we reduced to 20 dimensions in the first two and 10 dimensions in the second two cases (in these latter cases, we did not have enough neurons left to retain high dimensionality in the LEM space). For the study on LEM reductions after the removal of sensorimotor areas, we removed *n_max_* =max(#M1, #M2, #S1 units) from each area for each session. For underrepresented areas, we additionally discarded *n_max_ − n_area_* randomly chosen units. As before, given the lower number of neurons, we reduced to 10 dimensions. To investigate the dimensionality of the LEM space using the method of (*5*), we computed the average number of neighbors of all time points in the 20-dimensional LEM space in circles with increasing radii. The dimensionalities were then obtained as the slope of a line around the steepest point in a log–log plot of neighbors against radii.

For the dimensionality reduction with Isomap, we used Landmark–Isomap (*50*), which is more efficient for very large datasets. We set the number of neighbors to 0.5%, as for the LEM, and used 10% of the time points as landmarks. PCA reductions where computed on non-binarized spikes.

#### Behavioral labeling

We used the freely available tool MuViLab for the behavioral labeling of the videos. Two human annotators who were blinded to the neural data manually labeled the 48 sessions divided into 500-ms snippets. The 48 sessions were chosen based on them having a clear saddle-like shape and at least five significantly coupled units: Rat A—13 sessions recorded between 2017/06/08 and 2017/08/03, Rat B—16 sessions recorded between 2017/06/01 and 2017/08/21, Rat C—seven sessions between 2017/06/01 and 2017/06/29, Rat D—six sessions between 2017/06/08 and 2017/07/11, Rat E—three sessions between 2017/06/08 and 2017/06/22, and Rat F—three sessions between 2017/06/07 and 2017/06/30. The criteria for the behavioral classes were as follows: Step—the rat moved at least one paw but did not drink or rear at the same time; turn—the rat moved its head; drink—the rat drank from the spout or collected water drops from the mesh with its mouth; groom—the rat performed typical grooming movements; rear—the rat stood on its hind paws; rest—the rat showed no obvious movements. In rare cases, samples were excluded from labeling when the behavior of the rat was not visible because it was located near the borders of the arena. Examples of the different behaviors can be found at https://www.dropbox.com/sh/4uu3cmmmnnovqmb/AABWaTv9H_0MPgHOpx4tPOXwa?dl=0.

#### Single-unit behavioral coding

To establish the single-unit coding of a specific behavior or stimulus, it is common practice to compare the average firing rate of the unit prior to the event (baseline) and after it (response). In the case of self-initiated behaviors, however, it is difficult to unambiguously identify temporal windows that can be associated with a baseline or response. Thus, we tested whether a unit increased its firing rate during each of the six behavioral categories and compared this rate to the unit’s firing during the remainder of the recorded time. The test was performed using a Wilcoxon rank-sum test with Benjamini–Hochberg correction for multiple comparisons and *α* = 0.05.

In a second analysis, we aimed to compare the diversity in single-unit firing rates during two behaviors with the distance in the LEM space of the population vectors associated with such behaviors. To obtain the number of single-units that changed their firing rates during different behaviors, we divided the spike counts (500 ms binning) of each unit according to the six behavioral classes and performed a Kruskal–Wallis test. When the main effect was significant, we performed a post-hoc analysis to selectively compare the unit firing rates during each pair of behaviors. Significance was fixed at 0.05. Since the final aim of this analysis was to compare the average number of units that changed rates with the distance in the LEM space of the population vectors associated with different behaviors, we did not want the unequal sample size of the behavioral classes to affect the significance of the post-hoc tests. Therefore, before performing the Kruskal–Wallis test, we randomly selected an equal number of samples (equal to the sample size of the smallest class) from all behavioral classes for each unit. We then repeated the test 100 times and computed the average number (first across the 100 samplings and then across the session’s units) of significant post-hoc tests obtained for each class comparison and each session. Fig. S8a displays their average across sessions.

#### Similarities among behavioral representations across sessions

To investigate whether the relative positions of the neural representations associated with different behavioral classes were conserved across sessions and animals, we computed the Euclidean distance between the average behavioral population vectors of a session, then tested whether these distances were more similar to those observed in other sessions than to what would be expected by randomly shuffling the behavioral state labels. This was performed by first comparing the ranked distances between the polytope vertexes and then comparing the actual distance values. For each session *v*, we computed the Euclidean distance 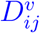 between the average population vector *ξ_i_* and *ξ_j_* of all pairs of behavioral classes *i* and *j*. Then, for each behavioral class *i*, we ranked the remaining classes *j* according to their distance 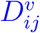 from *i*. For each pair of sessions *v* and *w* and each class *i*, we accounted for the similarities in ranked distances by defining the statistic 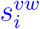 as the number of classes matching the same rank in the two sessions. For the six behavioral classes,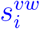 ranged between a maximum value of 5 (perfect match) to a minimum value of 0 (no match). The distribution of 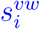 across all sessions was compared with a bootstrap distribution in which the same statistic, *s_boot_*, was computed over two random permutations of the numbers from 1 to 5. With six possible classes, there are 5! = 120 possible permutations of the remaining five classes, giving 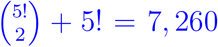 unordered pairs of random permutations. We thus used the Kolmogorov–Smirnov test to compare the distribution between the observed 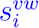 (*n* = 990 session pairs) and bootstrapped *s_boot_* (*n* = 7, 200) similarities.

The analysis described in the previous paragraph tested whether the distances between the pairs of behaviors (polytope vertexes) had a similar order (e.g., from the closest to the furthest) for different sessions or animals. To compare the actual distance values, we computed the matrix of pairwise Euclidean distances *D^v^* between the average class population vectors *ξ^i^* in the LEM space. Then, for each other session *w*, we performed a Procrustes transformation to rescale the behavioral population vectors of *w* with those of *v* and computed the distance matrix *D^w^* on the rescaled vectors. The Procrustes transformation did not affect the relative distance between vertexes but prevented differences in scale between the polytopes of different sessions from obscuring the quantity of interest. To quantify whether the set of relative distances between behavioral classes was, to some extent, maintained across sessions, we computed the difference between *D^v^* and *D^w^* as the Jeffries–Matusita distance 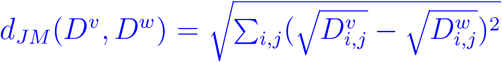, where *i* and *j* were indexes running over the six classes, and compared this difference with what we would obtain by chance. We employed the Jeffries–Matusita metric because it reduces the effect of outliers, but similar results were found with a Euclidean metric as well. The distribution 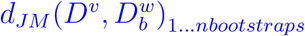 obtained with the original distance matrices was tested against the bootstrap distribution 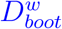 obtained by randomly permuting the behavioral labels associated with the population vectors of the session *w*. For each session pair (*v, w*), we then compared *d_JM_* (*D^v^, D^w^*) with those obtained on the bootstrapped *D^w^* and computed a p-value for the *H*_0_ of *d_JM_* (*D^v^, D^w^*) that was obtained by chance. The bootstrap sample included all possible class label permutations (*n* = 720). Figs. 3b and S10d show the significance of the comparison of each session pair when computed on all six behavioral classes, and Fig. S10e shows the same but with the “rest” class excluded.

#### Population-level decoding

We trained one deep neural network per session to classify the six behavioral classes given the 10-dimensional neural data in seven bins with 100 ms each as input. The data was min–max normalized (min and max were only calculated on training sets). The deep network architecture and training were almost identical to the network used for the decoding task above. However, we used only 200 units per layer and a dropout rate of 25%, and we chose a cross-validation strategy to deal with unbalanced classes. In the latter step, the available data was split into four parts of equal size. Four runs were conducted per session, using two parts as the training set, one as the validation set for early stopping, and the fourth as a test set. The final test results were calculated as the mean over all four test sets and runs. As for the decoding of the swing–stance status, we used weighted cross-entropy loss (more weight on less frequent classes) to deal with the class imbalance. All accuracies that we report were mean-per-class accuracies (balanced accuracies) to ensure that more frequent classes did not bias the results. While we used 10 dimensions for this behavioral decoding task—in line with the estimated dimensionality—only five dimensions remained for the area-specific dimensionality-reduced data since the lower number of neurons did not allow for a reduction in a higher-dimensional LEM space. For the supervised alignment procedure, we always restricted the analysis to four neural dimensions to avoid underdetermination. (That is, the remaining dimensions provided by LEM were not used – no completely new dimensionality reduction was computed.) We used Matlab’s *Procrustes* function to find a transformation between class means. Proper transformation was important because of the sign ambiguity of eigenvectors, which might otherwise have led to different orientations of the neural structures. Before alignment, both neural structures were normalized to the 0–1 range. An SVM with a Gaussian kernel (Matlab *fitcecoc*) was used as the classifier. Training was conducted with an equalized number of samples per class (i.e., the class with the fewest samples determined the number of samples taken from each class) and default parameters (kernel size 1). For the SVM classification, we did not use four-fold cross-validation as we did for the classification of neural networks (see above). Instead, we performed 20 repetitions with different samplings of the training set (Monte Carlo cross-validation).

**Figure S1:**
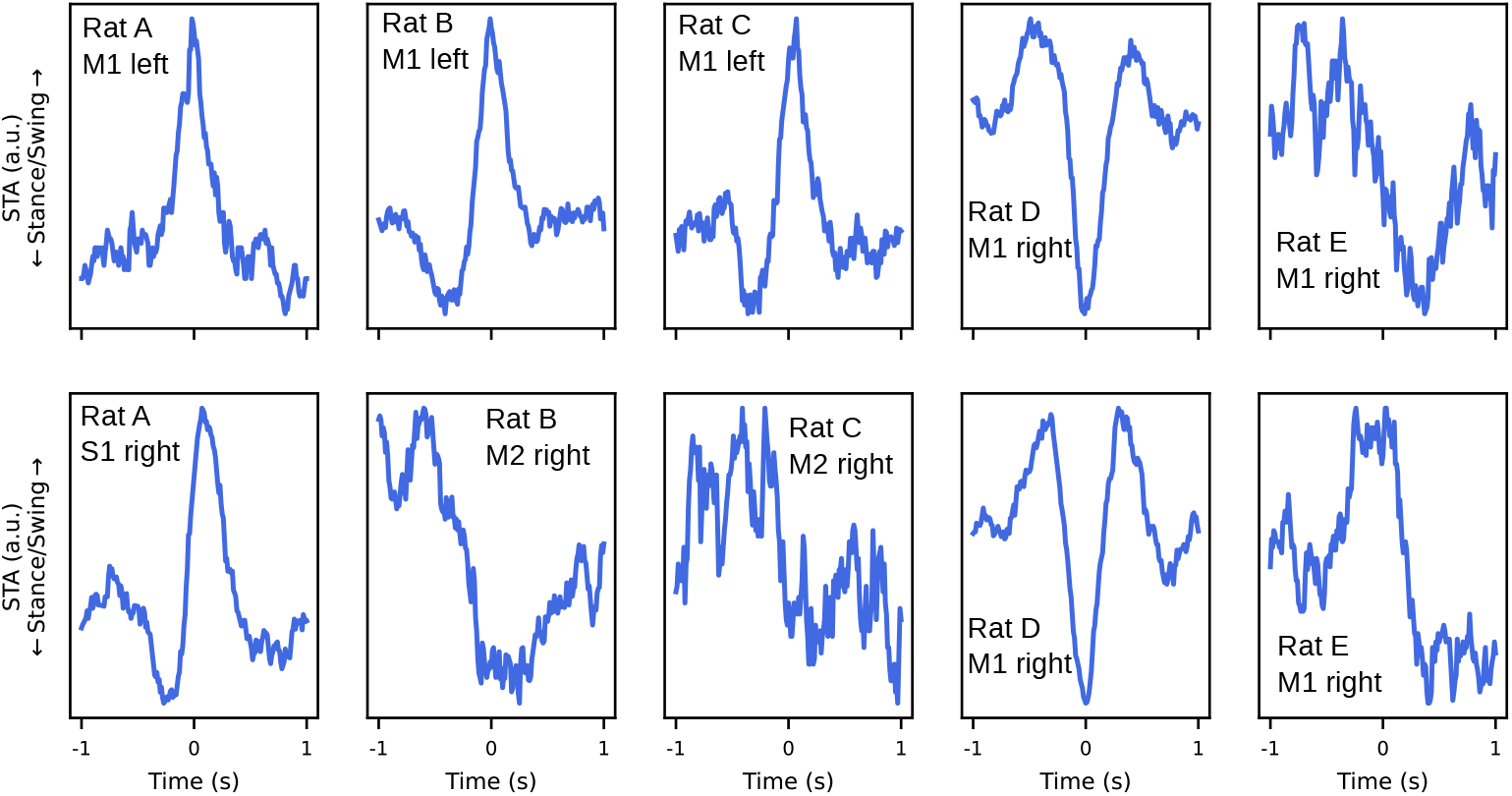
Significantly coupled neurons showed clear peaks in the STAPSSS. STAPSSS for the right front paw of 10 example neurons from different motor areas from five sessions of different rats. Extends Fig. 1d from the main paper.

**Table S1:** Statistics of the recording sessions. Dates of implantations and the recording periods for each animal.

|  | Rat A | Rat B | Rat C | Rat D | Rat E | Rat F |
| --- | --- | --- | --- | --- | --- | --- |
| Implantation | 20/04/2017 | 19/04/2017 | 27/04/2017 | 11/04/2017 | 25/04/2017 | 01/01/2017 |
| First recording | 01/06/2017 | 01/06/2017 | 01/06/2017 | 01/06/2017 | 01/06/2017 | 07/06/2017 |
| Last recording | 15/08/2017 | 21/08/2017 | 08/07/2017 | 21/08/2017 | 25/08/2017 | 22/08/2017 |

**Table S2:** ANOVA results for paw coupling. Paw coupling was defined as the ratio between the STAPSSS standard deviation and the control standard deviation (see main text). Three-way ANOVAs were calculated separately for each paw on all recorded neurons (*n* = 3, 723, main effects area, hemisphere, rat; interaction effect area and hemisphere). The table contains the corresponding *F* and *p* values.

| Paw | Area | Hemisphere | Area x Hemisphere | Rat |
| --- | --- | --- | --- | --- |
| Right front | 66.77, $p < 1e-28$ | 108.85, $p < 1e-24$ | 18.24, $p < 1e-07$ | 28.59, $p < 1e-27$ |
| Left front | 41.61, $p < 1e-17$ | 17.15, $p < 1e-4$ | 2.22, $p = .10$ | 37.82, $p < 1e-37$ |
| Right hind | 25.73, $p < 1e-11$ | 4.16, $p = .04$ | 5.63, $p = .003$ | 23.63, $p < 1e-22$ |
| Left hind | 67.47, $p < 1e-28$ | 6.38, $p = .01$ | 3.82, $p = .02$ | 13.52, $p < 1e-12$ |

**Figure S2:**
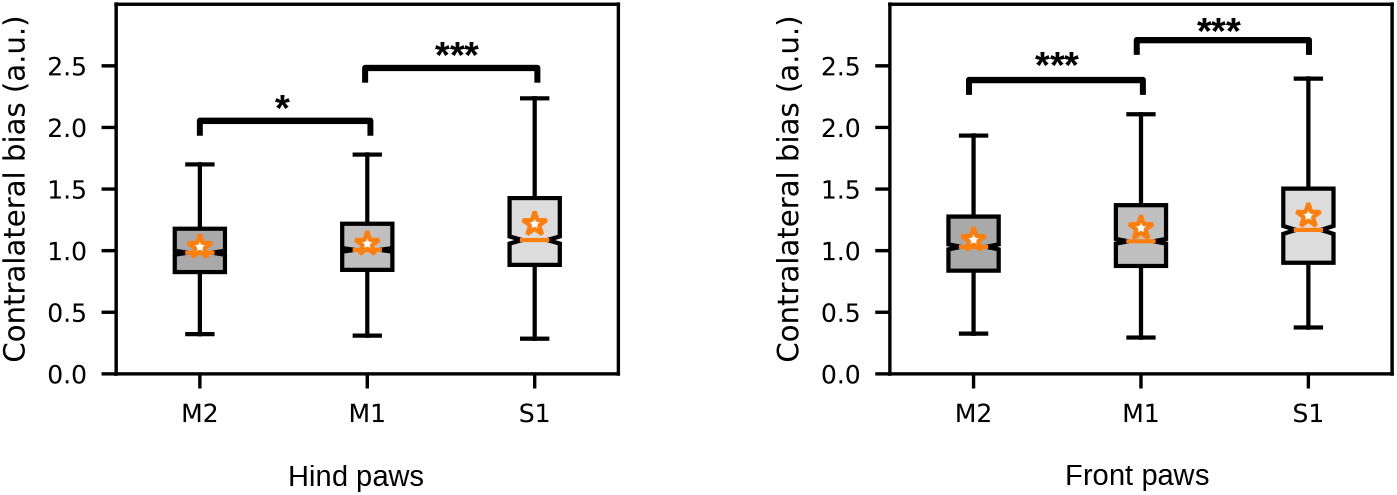
Contralateral bias was largest in S1. Contralateral bias for the front and hind paws per area, averaged over neurons. The bias increased from anterior to posterior regions for both the front and hind paws. Stars denote the results of the post-hoc Tukey–Kramer tests. Orange stars denote mean values, and notches denote the 95% confidence intervals for the median. See the main text for definitions of paw coupling and bias. \**p* < .05, \*\*\**p* < .001.

**Figure S3:**
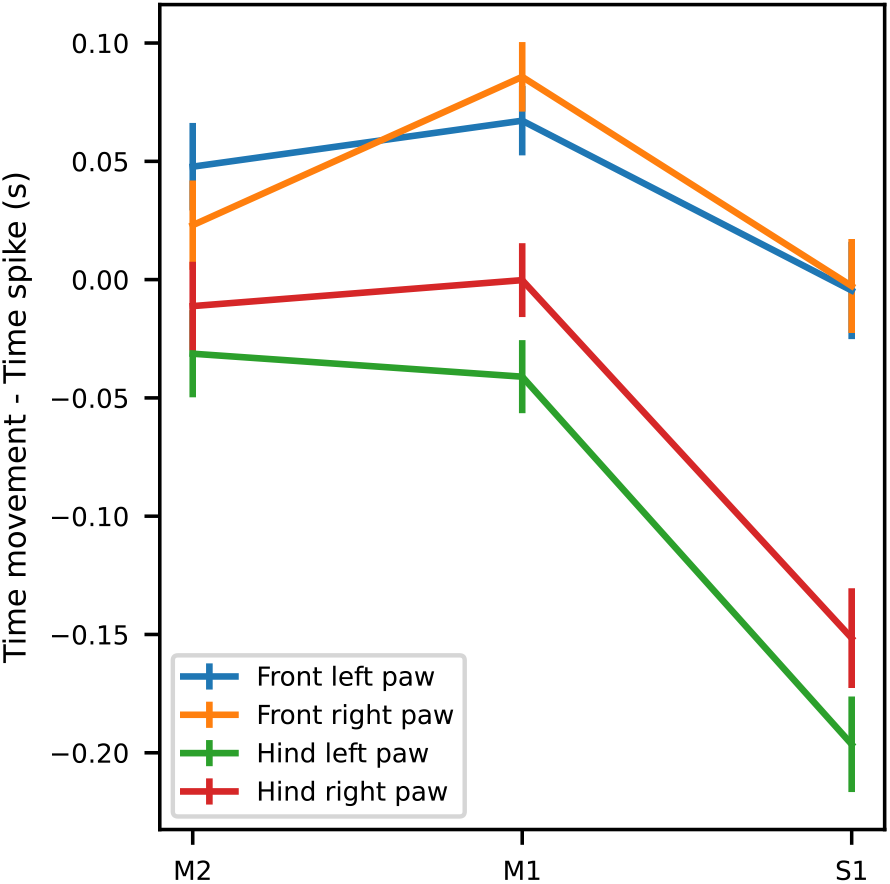
Temporal relationship between movement and brain-area-specific neuronal activity in M2, M1, and S1. Movement refers here to the STAPSSS peak. Negative values indicate that the spikes followed the movement (in the form of the STAPSSS peak); positive values indicate that the spikes preceded the movement. The spikes in S1 tended to occur after movements, significantly later than the spikes in M2 and M1. The mean and standard error of the mean over all neurons in each area are shown. Refers to the main paper’s Fig. 1.

**Figure S4:**
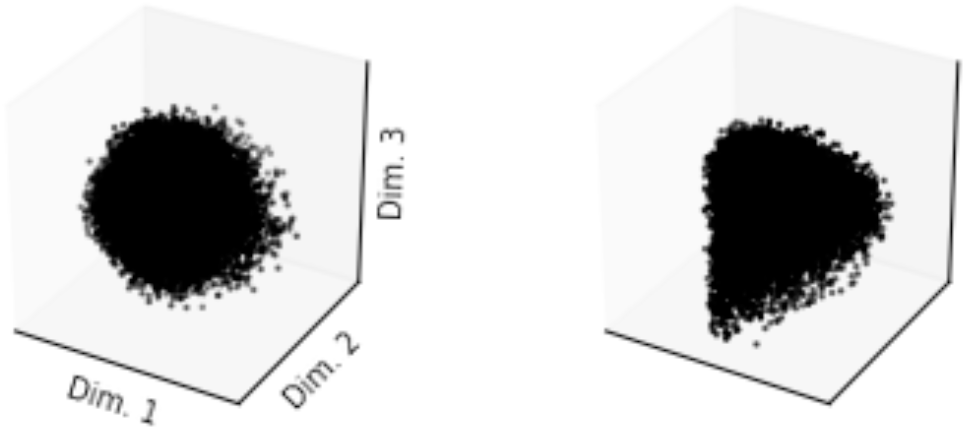
Some population structures did not show any apparent structure. Two example sessions (from Rats A and B, respectively) with random-like, low-dimensional neural projections. Refers to the main paper’s Fig. 2a.

**Figure S5:**
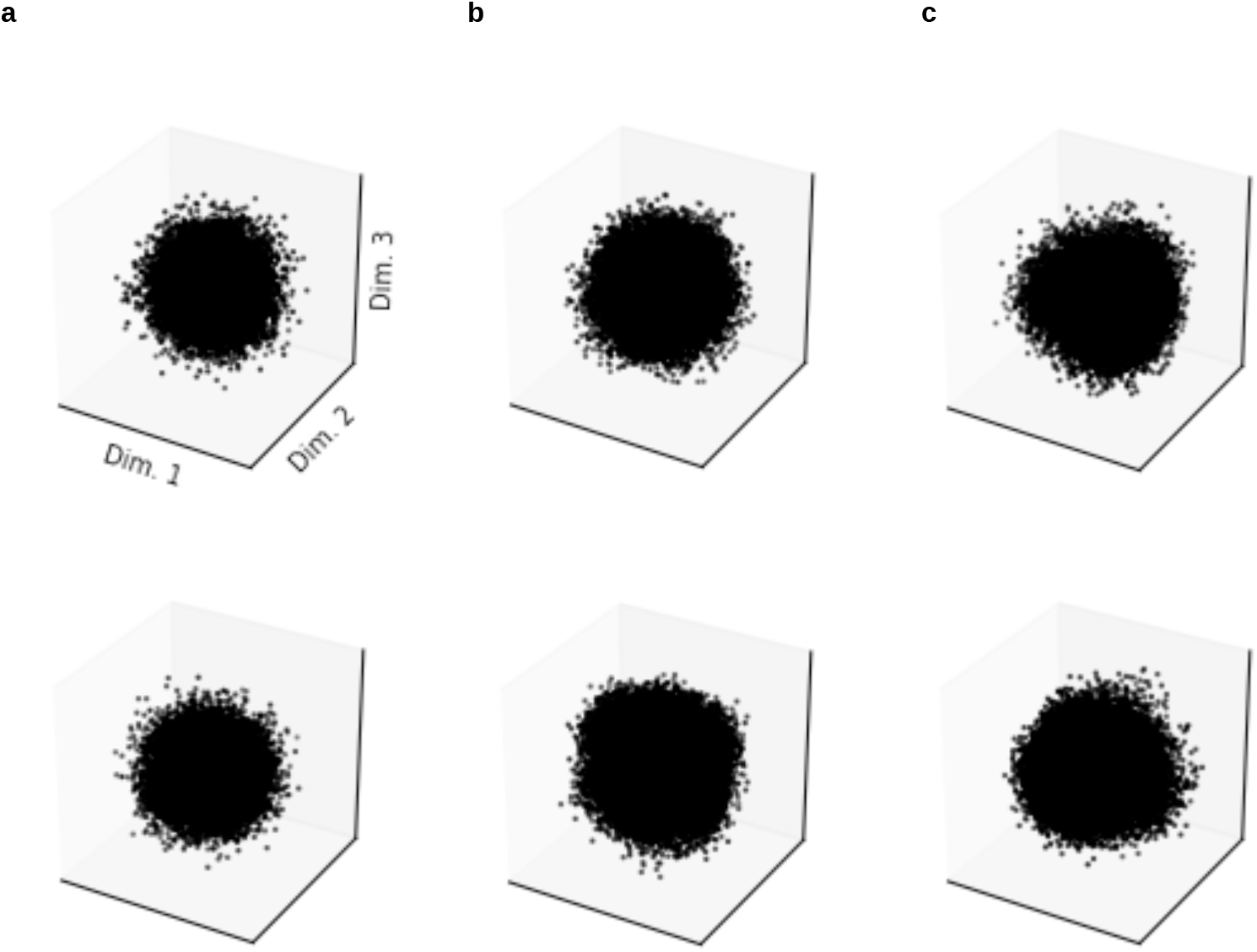
Control dimensionality reductions with shuffled neuronal activity did not show any apparent structure. LEM projections for neuron-shuffled (a), time-shuffled (b), and time-shifted (c) data for one session of Rat A (upper row) and Rat B (lower row). For neuron shuffling, units were permuted randomly for each time point. For time shuffling, time points were permuted randomly for each neuron. For time shifting, the spike trains of the neurons were randomly shifted against each other. The two sessions are the same as in Fig. 2b of the main paper.

**Figure S6:**
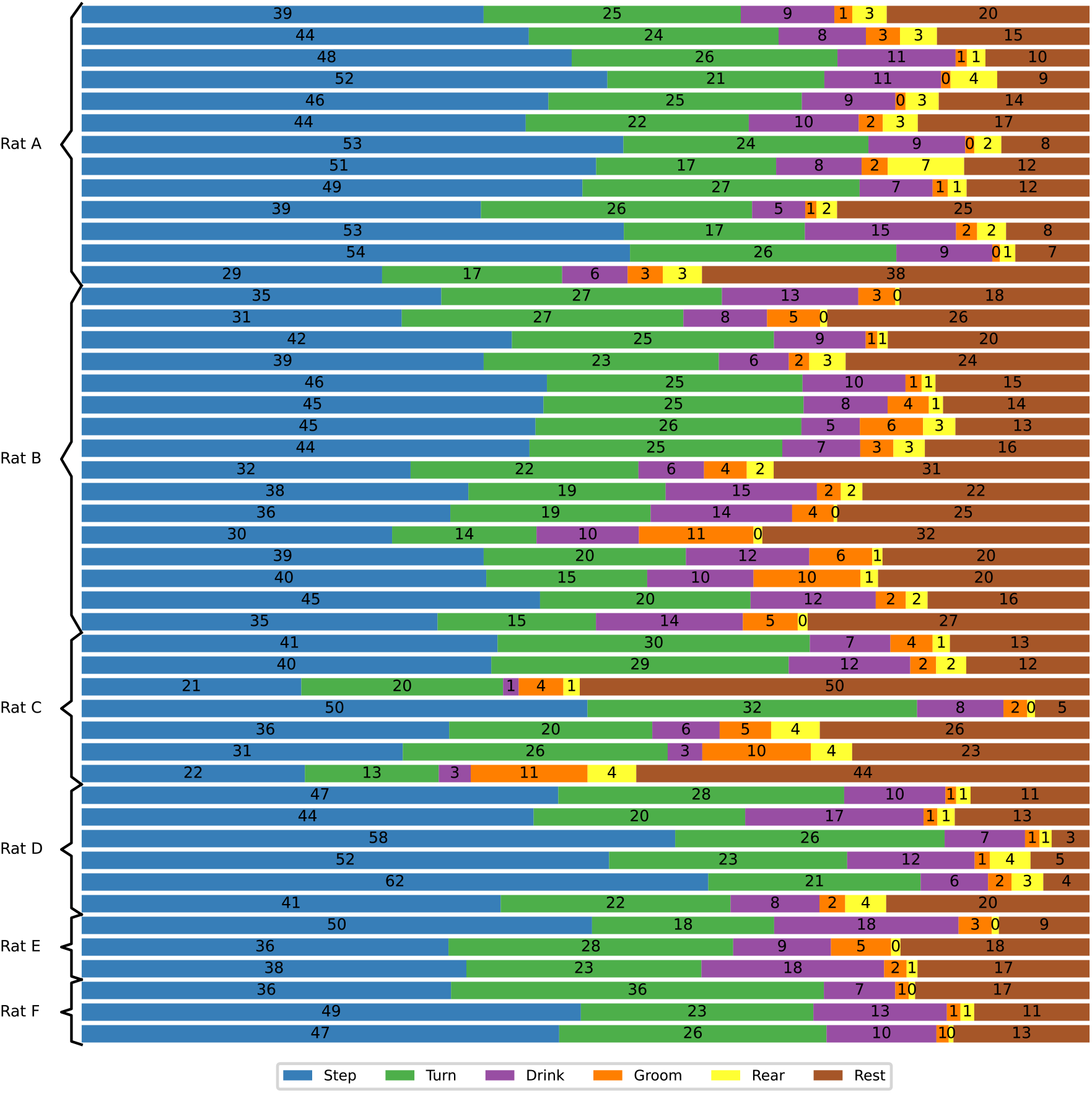
Histograms of behavior illustrating the distribution of behavioral classes. One row reflects one session. Provides background for Fig. 2 in the main paper.

**Figure S7:**
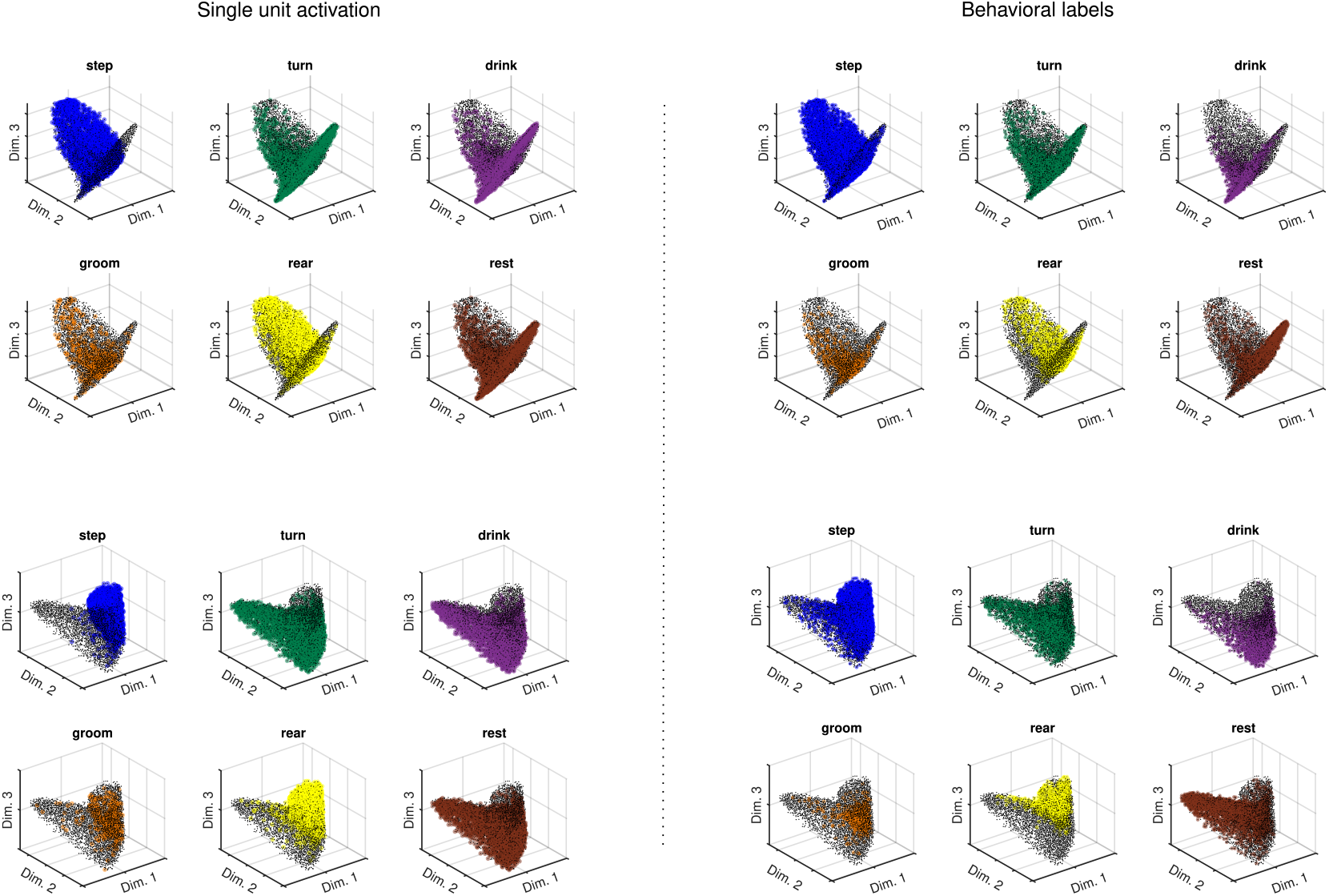
Comparison of single-unit firing patterns and localization of behaviors in the LEM space. (Left) The LEM manifold from one example session of Rat A is color-coded according to the firing of six different example single-units significantly responding to the six behavioral classes. Colored dots mark time points when the unit fired above its 75th empirical quartile, while black dots mark any other time point. (Right) On the same LEM manifold as in the left panel, colored dots mark time points corresponding to the six behavioral labels. Different orientations of the same manifold are shown from top to bottom.

**Figure S8:**
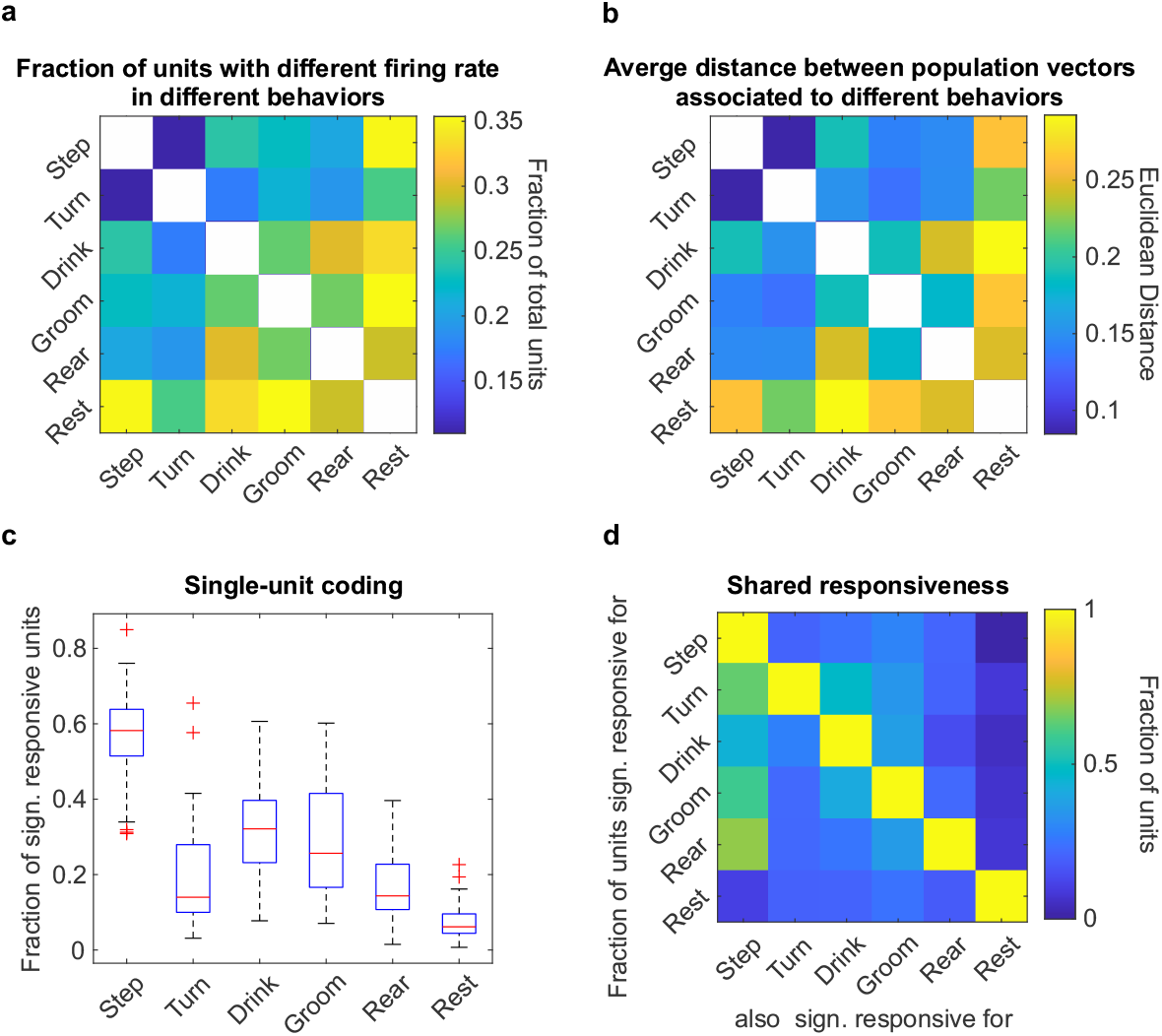
Single-unit coding of different behaviors. (a) Fraction of single-units changing their average firing rate when active during different behaviors as the fraction of significant post-hoc comparisons (*α* = 0.05) of a Kruskal–Wallis test on the firing rate of single-units during the six identified behaviors. Differences in sample size across classes were compensated with down-sampling (see Methods). (b) Average distance between the population vectors of the LEM space (dim = 10) associated with different behaviors. (c) Fraction of units significantly more active during each of the behaviors (Wilcoxon rank-sum test, with Benjamini–Hochberg correction for multiple comparisons, *α* = 0.05). The median (red line) across sessions, the 25th and 75th percentiles (blue), the most extreme data points (whiskers), and outliers (crosses) are shown. (d) Based on (c), the fraction of units with shared responsiveness to multiple behaviors.

**Figure S9:**
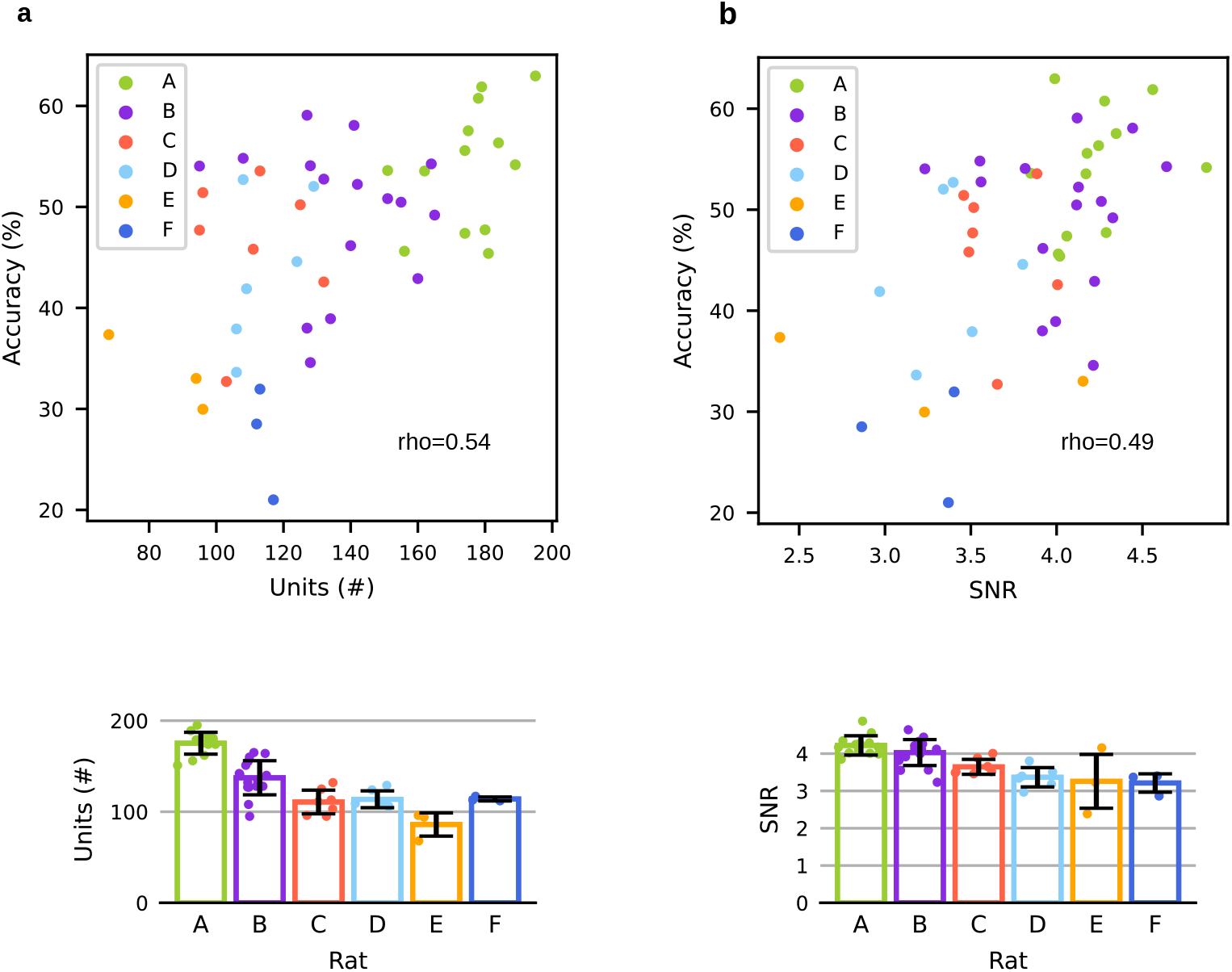
Correlation between accuracies and units/SNR. (a) Top: Accuracies versus the number of units per session for the six rats. Bottom: Average number of units per rat and error bars for the standard deviation across sessions. (b) Top: Accuracies versus the mean SNR per session for the six rats. Bottom: Average SNR per rat, with error bars for the standard deviation. Refers to Fig. 2.

**Figure S10:**
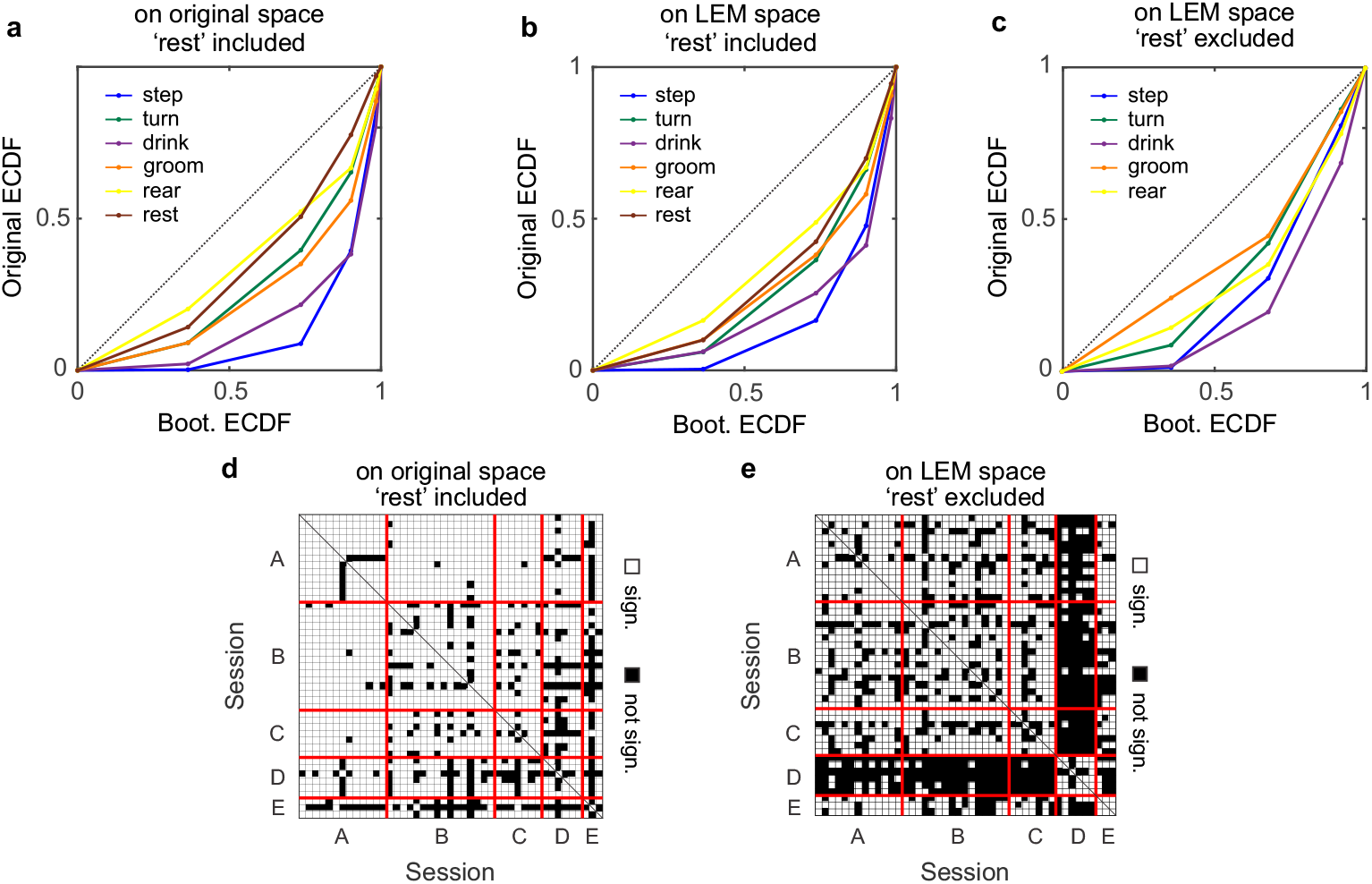
Similarity among the polytopes of different sessions. (a–c) Probability– probability (p–p) plot comparing the original and bootstrapped empirical cumulative distribution function (ECDF) of the statistic *s^vw^*, which compares the ranked distances between the polytope vertexes across sessions (see Methods for a formal definition). The ECDFs of *s^vw^* were computed for each behavioral class *i* (color-coded) on the original recordings (a), on the 20-dimensional LEM space (b), and on the 20-dimensional LEM space with the class “rest” excluded from the test (c). In a p–p plot, equal distributions overlap with the diagonal (dotted line). (d–e) are the same as in Fig. 3 b, but with computing distances on the original recording space (d) and on the 20-dimensional LEM space with the “rest” class excluded from the test (e). Of the 990 possible session pairs, 84% and 61% in (d) and (e), respectively, had a p-value below 0.05.

**Figure S11:**
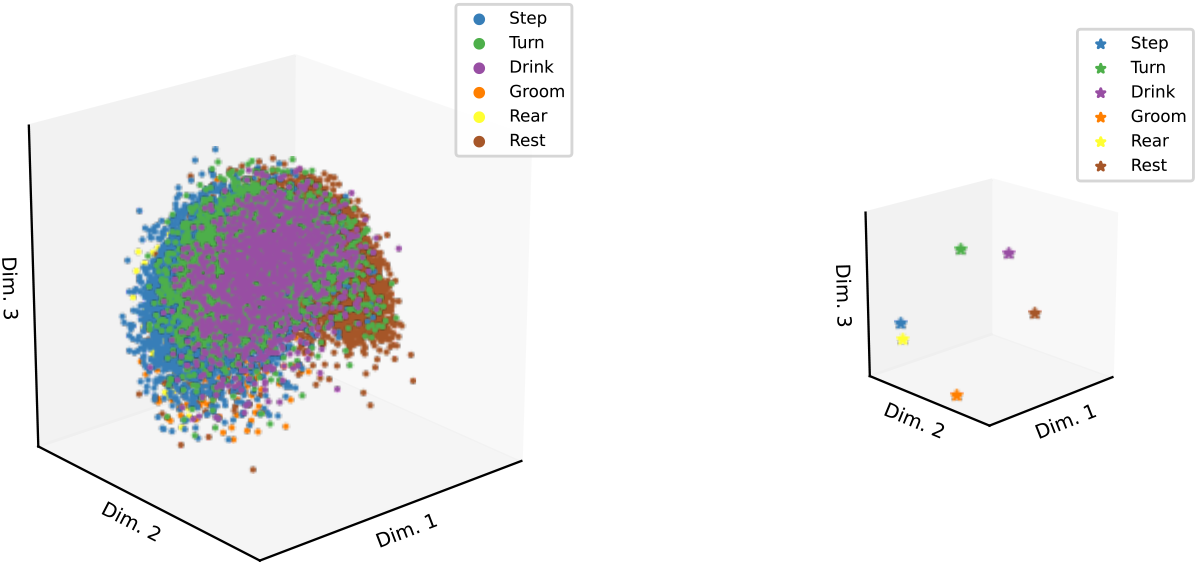
Population structure in the Isomap space. Left: All points. Right: For better visualization, only the averages of the six behavioral classes were plotted. One session of Rat A is shown here.

**Figure S12:**
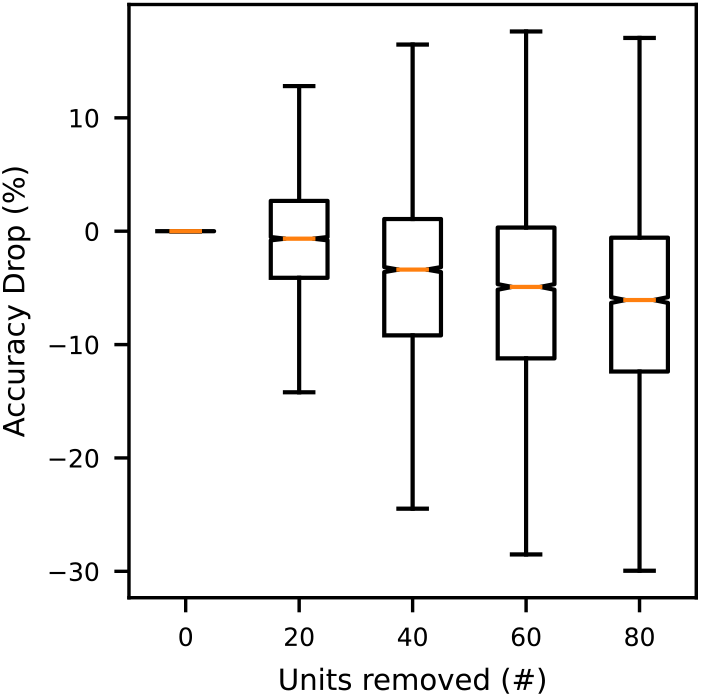
Generalization worsened with fewer units. We repeated the generalization experiment from Fig. 5a for all sessions with a generalization accuracy of at least 55% (19 sessions). The accuracy decreased for LEM structures that were computed after removing 20, 40, 60, or 80 units from each session compared to the accuracies with the full number of units. Thus, accuracy decreased with fewer units.

**Figure S13:**
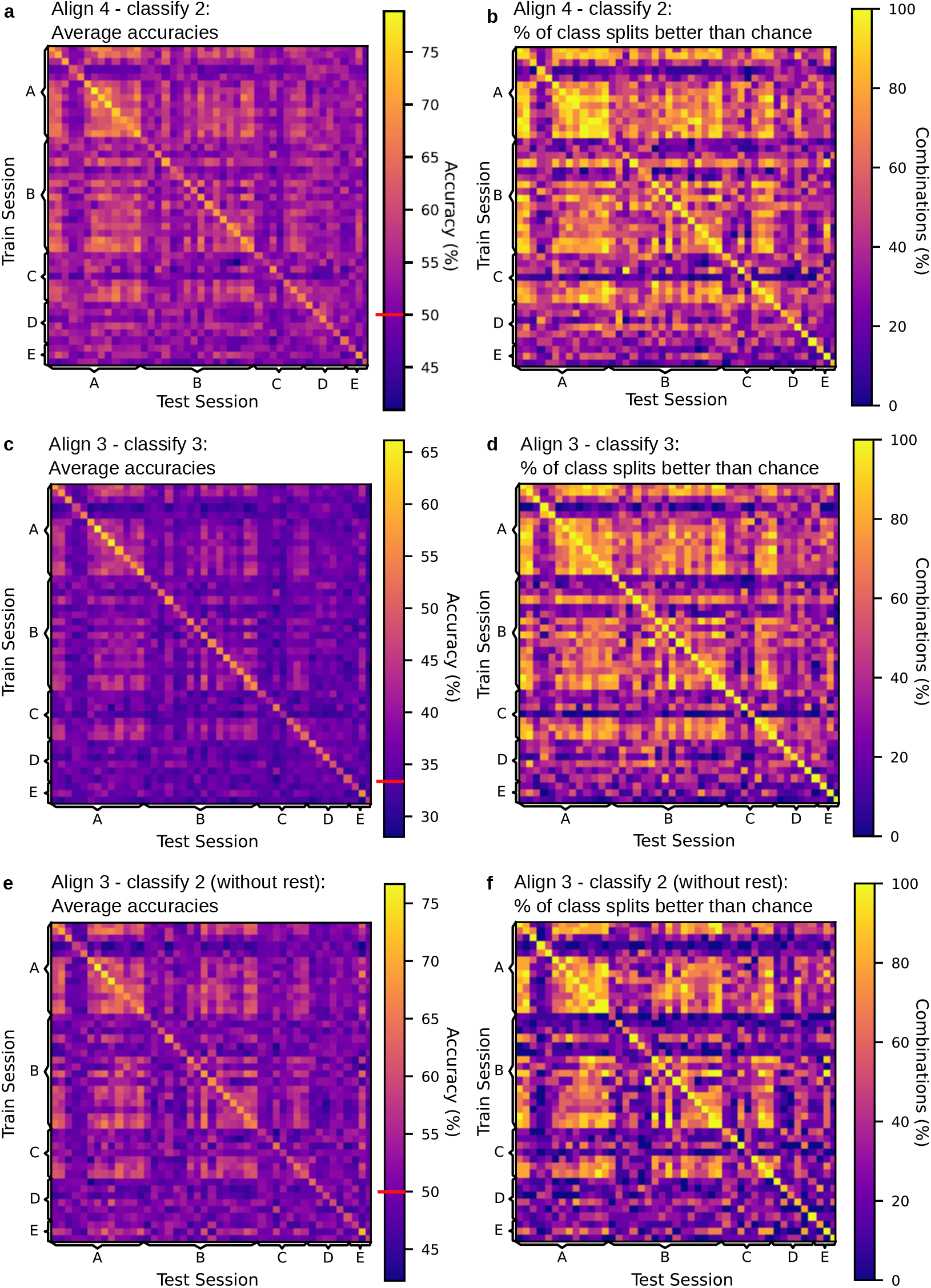
Further cross-subject and cross-session generalization studies. Further cross-subject generalization experiments and performance above chance. For all plots, training and test data on the diagonal originated from the same session. Off-diagonal entries show testing on data other than the training session. (a) Mean per-class accuracies across training and test sessions when aligning on four and testing on two classes. The chance level was 50% (red line). Values were averaged over 20 runs and 15 possible splits of the six behavioral classes into alignment and decoding sets. We used the same plot as in Fig. 5a in the main paper. (b) Percentage of splits of the six behavioral classes into the alignment and decoding sets (out of 15 splits) that were classified with a significantly higher mean per-class decoding accuracy than chance (50%). Significance was calculated over 20 training runs at a .05 significance level with Bonferroni correction using a one-tailed sign test. In total, the accuracy was significantly better than chance in 47.55%(14, 445*/*(15 *∗* 45 *∗* 45) = 14, 445*/*30, 375) of the experiments. This figure refers to the main paper’s Fig. 5a. (c–f) We conducted further generalization experiments with a more difficult setting (align on three classes and classify three classes, c–d; experiment without the “rest” class, e–f). (c) Mean per-class accuracies across training and test sessions when aligning on three and testing on three classes. The chance level was 33.33%. Values were averaged over 20 runs and 20 possible splits of the six behavioral classes into alignment and decoding sets. (d) Percentage of splits with above-chance per-class decoding accuracy, as in (b), but for the experiment that aligns on three classes and tests with three classes, with 20 combinations in total and a chance level of 33.33% (red line). In total, in 44.28%(17, 934*/*40, 500) of the experiments, the accuracy was significantly better than chance. (e) Mean per-class accuracies across training and test sessions when aligning on three and testing on two classes, without the “rest” class. The chance level was 50% (red line). Values were averaged over 20 runs and 10 possible splits of the six behavioral classes into alignment/decoding sets. (f) This is the same as in (b) and (d) for aligning on three and testing on two classes without the class “rest,” with 10 combinations in total and a chance level of 50%. In total, in 37.25%(7, 545*/*20, 250) of the experiments, the accuracy was significantly better than chance. Extends Fig. 5 in the main paper.

**Figure S14:**
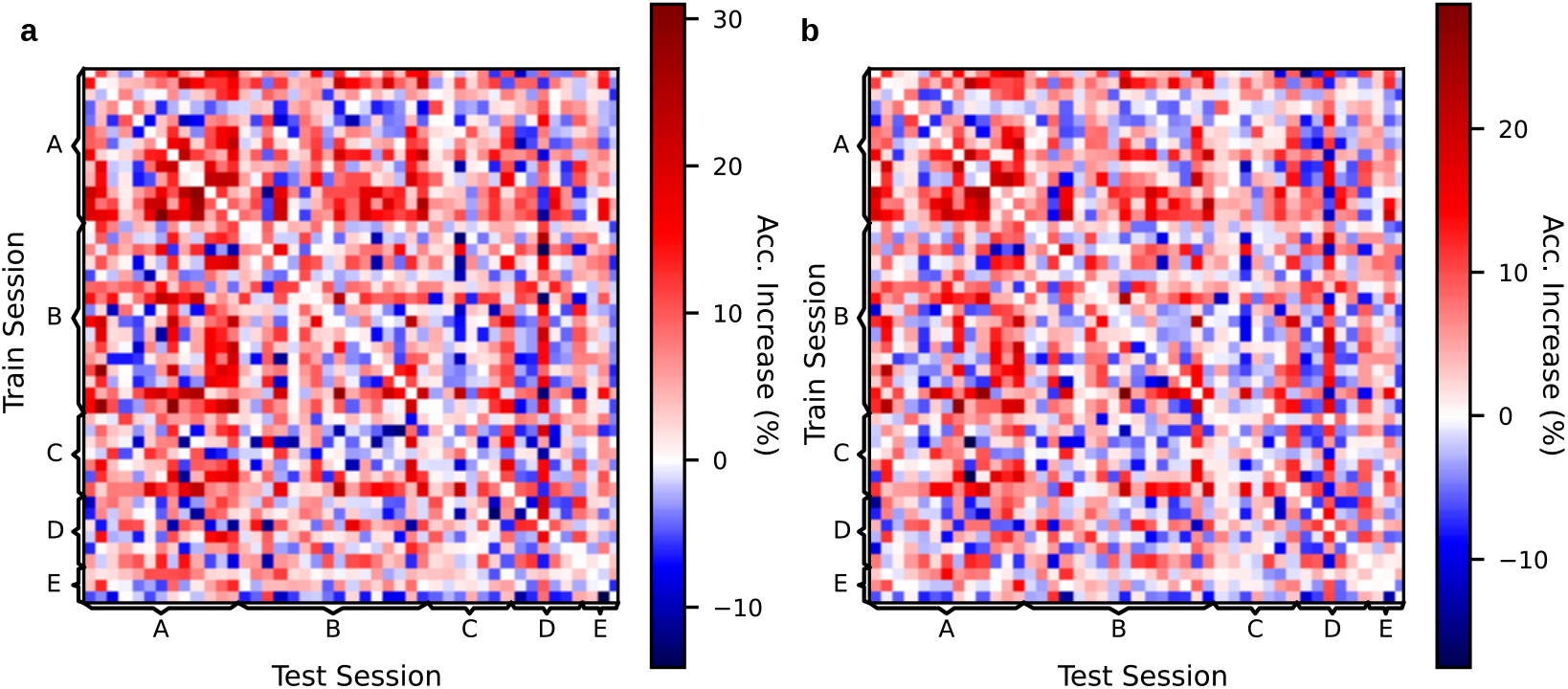
Decoding accuracy gain through neural manifold alignment. Accuracy gains for aligned versus unaligned neural structures for the decoding of (a) two classes and (b) three classes. Values are averaged over all class combinations. In most cases, the accuracies were higher after alignment (red color spectrum) by up to 20–30%. Refers to Fig. 5a in the main paper and Fig. S13a–d.

**Figure S15:**
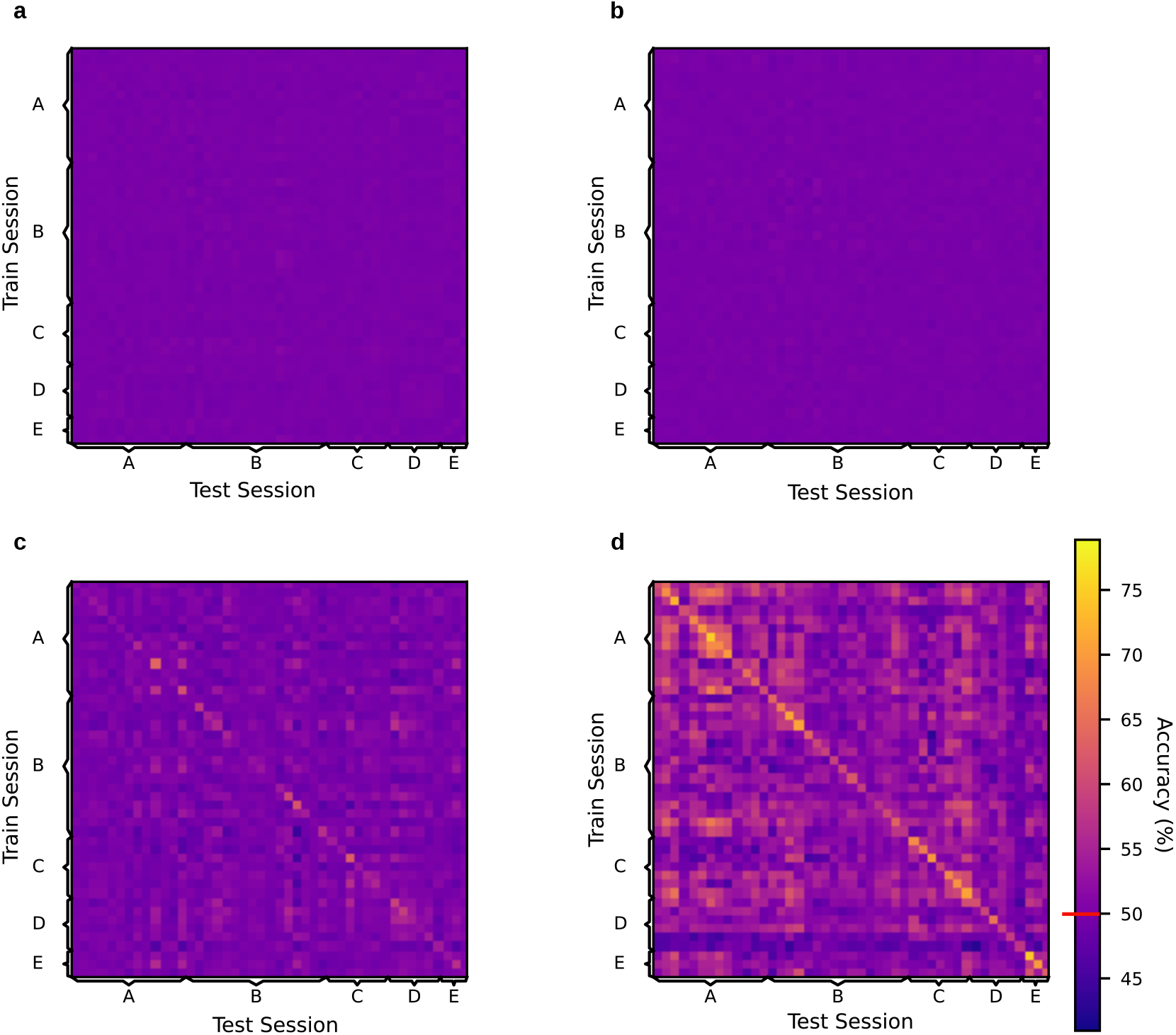
Control generalization experiments. Generalization results on neuron-shuffled
(a), time-shuffled (b), and time-shifted (c) data, as well as the LEM space from non-binarized
spikes (d). Mean per-class accuracies across training and test sessions when aligning on four
and testing on two classes (chance level of 50%, red line) are shown, as in Fig. 5a.

## Footnotes

1 Quotient of recorded cells and estimated total number of cells (approximated for the area covered by the implanted electrodes by assuming a cortical thickness of 2mm and a density of 90k neurons per mm^3^ (*26*)).

## References

1. G. R. Yang, M. R. Joglekar, H. F. Song, W. T. Newsome, and X.-J. Wang, “Task representations in neural networks trained to perform many cognitive tasks,” Nature neuroscience, vol. 22, no. 2, pp. 297–306, 2019.

2. S. Vyas, M. D. Golub, D. Sussillo, and K. V. Shenoy, “Computation through neural population dynamics,” Annual Review of Neuroscience, vol. 43, pp. 249–275, 2020.

3. S. D. Stavisky, F. R. Willett, G. H. Wilson, B. A. Murphy, P. Rezaii, D. T. Avansino, W. D. Memberg, J. P. Miller, R. F. Kirsch, L. R. Hochberg, et al., “Neural ensemble dynamics in dorsal motor cortex during speech in people with paralysis,” Elife, vol. 8, p. e46015, 2019.

4. M. G. Perich, J. A. Gallego, and L. E. Miller, “A neural population mechanism for rapid learning,” Neuron, vol. 100, no. 4, pp. 964–976, 2018.

5. A. Rubin, L. Sheintuch, N. Brande-Eilat, O. Pinchasof, Y. Rechavi, N. Geva, and Y. Ziv, “Revealing neural correlates of behavior without behavioral measurements,” Nature communications, vol. 10, no. 1, pp. 1–14, 2019.

6. M. Rabinovich, R. Huerta, and G. Laurent, “Transient dynamics for neural processing,” Science, vol. 321, no. 5885, pp. 48–50, 2008.

7. K. V. Shenoy, M. Sahani, and M. M. Churchland, “Cortical control of arm movements: a dynamical systems perspective,” Annual review of neuroscience, vol. 36, 2013.

8. J. A. Michaels, B. Dann, and H. Scherberger, “Neural population dynamics during reaching are better explained by a dynamical system than representational tuning,” PLoS computational biology, vol. 12, no. 11, p. e1005175, 2016.

9. D. V. Buonomano and W. Maass, “State-dependent computations: spatiotemporal processing in cortical networks,” Nature Reviews Neuroscience, vol. 10, no. 2, pp. 113–125, 2009.

10. E. E. Fetz, “Are movement parameters recognizably coded in the activity of single neurons?,” Behavioral and Brain Sciences, vol. 15, no. 4, pp. 679–690, 1992.

11. M. M. Churchland, J. P. Cunningham, M. T. Kaufman, J. D. Foster, P. Nuyujukian, S. I. Ryu, and K. V. Shenoy, “Neural population dynamics during reaching,” Nature, vol. 487, no. 7405, p. 51, 2012.

12. M. T. Kaufman, M. M. Churchland, S. I. Ryu, and K. V. Shenoy, “Cortical activity in the null space: permitting preparation without movement,” Nature neuroscience, vol. 17, no. 3, p. 440, 2014.

13. P. T. Sadtler, K. M. Quick, M. D. Golub, S. M. Chase, S. I. Ryu, E. C. Tyler-Kabara, M. Y. Byron, and A. P. Batista, “Neural constraints on learning,” Nature, vol. 512, no. 7515, p. 423, 2014.

14. J. A. Gallego, M. G. Perich, L. E. Miller, and S. A. Solla, “Neural manifolds for the control of movement,” Neuron, vol. 94, no. 5, pp. 978–984, 2017.

15. A. Miri, C. L. Warriner, J. S. Seely, G. F. Elsayed, J. P. Cunningham, M. M. Churchland, and T. M. Jessell, “Behaviorally selective engagement of short-latency effector pathways by motor cortex,” Neuron, vol. 95, no. 3, pp. 683–696, 2017.

16. J. A. Gallego, M. G. Perich, S. N. Naufel, C. Ethier, S. A. Solla, and L. E. Miller, “Cortical population activity within a preserved neural manifold underlies multiple motor behaviors,” Nature communications, vol. 9, no. 1, p. 4233, 2018.

17. J. A. Hennig, M. D. Golub, P. J. Lund, P. T. Sadtler, E. R. Oby, K. M. Quick, S. I. Ryu, E. C. Tyler-Kabara, A. P. Batista, M. Y. Byron, et al., “Constraints on neural redundancy,” Elife, vol. 7, p. e36774, 2018.

18. A. A. Russo, S. R. Bittner, S. M. Perkins, J. S. Seely, B. M. London, A. H. Lara, A. Miri, N. J. Marshall, A. Kohn, T. M. Jessell, et al., “Motor cortex embeds muscle-like commands in an untangled population response,” Neuron, vol. 97, no. 4, pp. 953–966, 2018.

19. C. Pandarinath, K. C. Ames, A. A. Russo, A. Farshchian, L. E. Miller, E. L. Dyer, and J. C. Kao, “Latent factors and dynamics in motor cortex and their application to brain–machine interfaces,” Journal of Neuroscience, vol. 38, no. 44, pp. 9390–9401, 2018.

20. G. F. Elsayed, A. H. Lara, M. T. Kaufman, M. M. Churchland, and J. P. Cunningham, “Reorganization between preparatory and movement population responses in motor cortex,” Nature communications, vol. 7, p. 13239, 2016.

21. C. Pandarinath, D. J. O’Shea, J. Collins, R. Jozefowicz, S. D. Stavisky, J. C. Kao, E. M. Trautmann, M. T. Kaufman, S. I. Ryu, L. R. Hochberg, et al., “Inferring single-trial neural population dynamics using sequential auto-encoders,” Nature methods, vol. 15, no. 10, pp. 805–815, 2018.

22. M. Belkin and P. Niyogi, “Laplacian eigenmaps for dimensionality reduction and data representation,” Neural computation, vol. 15, no. 6, pp. 1373–1396, 2003.

23. S. Musall, M. T. Kaufman, A. L. Juavinett, S. Gluf, and A. K. Churchland, “Single-trial neural dynamics are dominated by richly varied movements,” Nature neuroscience, vol. 22, no. 10, pp. 1677–1686, 2019.

24. G. Paxinos and C. Watson, The rat brain in stereotaxic coordinates: hard cover edition. Elsevier, 2006.

25. P. A. Kells, S. H. Gautam, L. Fakhraei, J. Li, and W. L. Shew, “Strong neuron-to-body coupling implies weak neuron-to-neuron coupling in motor cortex,” Nature communications, vol. 10, no. 1, p. 1575, 2019.

26. V. Braitenberg and A. Schüz, Cortex: statistics and geometry of neuronal connectivity. Springer Science & Business Media, 2013.

27. J. V. Haxby, J. S. Guntupalli, A. C. Connolly, Y. O. Halchenko, B. R. Conroy, M. I. Gobbini, M. Hanke, and P. J. Ramadge, “A common, high-dimensional model of the representational space in human ventral temporal cortex,” Neuron, vol. 72, no. 2, pp. 404–416, 2011.

28. H.-T. Chen, J. R. Manning, and M. A. van der Meer, “Between-subject prediction reveals a shared representational geometry in the rodent hippocampus,” bioRxiv, 2020.

29. J. DiGiovanna, N. Dominici, L. Friedli, J. Rigosa, S. Duis, J. Kreider, J. Beauparlant, R. Van Den Brand, M. Schieppati, S. Micera, et al., “Engagement of the rat hindlimb motor cortex across natural locomotor behaviors,” Journal of Neuroscience, vol. 36, no. 40, pp. 10440– 10455, 2016.

30. S. Soma, A. Saiki, J. Yoshida, A. Rìos, M. Kawabata, Y. Sakai, and Y. Isomura, “Distinct laterality in forelimb-movement representations of rat primary and secondary motor cortical neurons with intratelencephalic and pyramidal tract projections,” Journal of Neuroscience, vol. 37, no. 45, pp. 10904–10916, 2017.

31. A. A. Russo, R. Khajeh, S. R. Bittner, S. M. Perkins, J. P. Cunningham, L. F. Abbott, and M. M. Churchland, “Neural trajectories in the supplementary motor area and motor cortex exhibit distinct geometries, compatible with different classes of computation,” Neuron, vol. 107, no. 4, pp. 745–758, 2020.

32. C. J. MacDowell and T. J. Buschman, “Low-dimensional spatio-temporal dynamics underlie cortex wide neural activity,” bioRxiv, 2020.

33. P. Gao and S. Ganguli, “On simplicity and complexity in the brave new world of large-scale neuroscience,” Current opinion in neurobiology, vol. 32, pp. 148–155, 2015.

34. J. P. Cunningham and M. Y. Byron, “Dimensionality reduction for large-scale neural recordings,” Nature neuroscience, vol. 17, no. 11, p. 1500, 2014.

35. J. A. Gallego, M. G. Perich, R. H. Chowdhury, S. A. Solla, and L. E. Miller, “Long-term stability of cortical population dynamics underlying consistent behavior,” Nature Neuroscience, pp. 1–11, 2020.

36. B. Mimica, B. A. Dunn, T. Tombaz, V. S. Bojja, and J. R. Whitlock, “Efficient cortical coding of 3d posture in freely behaving rats,” Science, vol. 362, no. 6414, pp. 584–589, 2018.

37. N. Kriegeskorte, M. Mur, and P. A. Bandettini, “Representational similarity analysis-connecting the branches of systems neuroscience,” Frontiers in systems neuroscience, vol. 2, p. 4, 2008.

38. J. V. Haxby, A. C. Connolly, and J. S. Guntupalli, “Decoding neural representational spaces using multivariate pattern analysis,” Annual review of neuroscience, vol. 37, pp. 435–456, 2014.

39. U. Hasson, Y. Nir, I. Levy, G. Fuhrmann, and R. Malach, “Intersubject synchronization of cortical activity during natural vision,” science, vol. 303, no. 5664, pp. 1634–1640, 2004.

40. J. S. Guntupalli, M. Hanke, Y. O. Halchenko, A. C. Connolly, P. J. Ramadge, and J. V. Haxby, “A model of representational spaces in human cortex,” Cerebral cortex, vol. 26, no. 6, pp. 2919–2934, 2016.

41. J. Chen, Y. C. Leong, C. J. Honey, C. H. Yong, K. A. Norman, and U. Hasson, “Shared memories reveal shared structure in neural activity across individuals,” Nature neuroscience, vol. 20, no. 1, pp. 115–125, 2017.

42. G. Zhang, V. Davoodnia, A. Sepas-Moghaddam, Y. Zhang, and A. Etemad, “Classification of hand movements from eeg using a deep attention-based lstm network,” IEEE Sensors Journal, vol. 20, no. 6, pp. 3113–3122, 2019.

43. N. Rabin, M. Kahlon, S. Malayev, and A. Ratnovsky, “Classification of human hand movements based on emg signals using nonlinear dimensionality reduction and data fusion techniques,” Expert Systems with Applications, vol. 149, p. 113281, 2020.

44. J. Zhou, C. Jia, M. Montesinos-Cartagena, M. P. Gardner, W. Zong, and G. Schoenbaum, “Evolving schema representations in orbitofrontal ensembles during learning,” Nature, vol. 590, no. 7847, pp. 606–611, 2021.

45. H.-T. Chen, J. R. Manning, and M. A. van der Meer, “Between-subject prediction reveals a shared representational geometry in the rodent hippocampus,” Current Biology, vol. 31, no. 19, pp. 4293–4304.e5, 2021.

46. W. E. Allen, M. Z. Chen, N. Pichamoorthy, R. H. Tien, M. Pachitariu, L. Luo, and K. Deisseroth, “Thirst regulates motivated behavior through modulation of brainwide neural population dynamics,” Science, vol. 364, no. 6437, pp. 253–253, 2019.

47. D. P. Kingma and J. Ba, “Adam: A method for stochastic optimization,” arXiv preprint arXiv:1412.6980, 2014.

48. U. Von Luxburg, “A tutorial on spectral clustering,” Statistics and computing, vol. 17, no. 4, pp. 395–416, 2007.

49. D. Eriksson, M. Heiland, A. Schneider, and I. Diester, “Distinct dynamics of neuronal activity during concurrent motor planning and execution,” Nature communications, vol. 12, no. 1, pp. 1–18, 2021.

50. V. De Silva and J. B. Tenenbaum, “Global versus local methods in nonlinear dimensionality reduction.,” in NIPS, vol. 15, pp. 705–712, 2002.

